# Minimization of a harmful cross-talk between mitotic checkpoint silencing and error correction

**DOI:** 10.1101/459594

**Authors:** Babhrubahan Roy, Vikash Verma, Janice Sim, Adrienne Fontan, Ajit P. Joglekar

**Affiliations:** Cell & Developmental Biology, University of Michigan Medical School, Ann Arbor; Biophysics, University of Michigan, Ann Arbor

## Abstract

Accurate chromosome segregation during cell division requires that the pair of sister kinetochores on each chromosome attach to microtubules originating from opposite spindle poles. This is ensured by the combined action of the Spindle Assembly Checkpoint (SAC), which detects unattached kinetochores, and an error correction mechanism that destabilizes incorrect attachment of both sister kinetochores to the same spindle pole. These processes are downregulated by Protein Phosphatase 1 (PP1), which both silences the SAC and stabilizes kinetochore-microtubule attachments. We find that this dual PP1 role can be problematic: if PP1 is recruited to the kinetochore for SAC silencing prior to chromosome biorientation, it interferes with error correction. We show that to mitigate this cross-talk, the yeast kinetochore uses independent PP1 sources to stabilize correct attachments and to silence the SAC, and also delays the recruitment of PP1 for SAC silencing. Consequently, chromosome biorientation precedes SAC silencing ensuring accurate chromosome segregation.

## Introduction

During cell division, chromosomes often form syntelic attachments wherein both sister kinetochores establish end-on attachments with microtubules from the same spindle pole (Fig. 1A). For accurate chromosome segregation, these erroneous attachments must be corrected before the cell enters anaphase. However, recent studies show that end-on kinetochore-microtubule attachments, whether they are monopolar, syntelic, or bipolar, can silence the SAC (Etemad et al., 2015; Tauchman et al., 2015). Therefore, to minimize chromosome missegregation, the kinetochore must allow SAC silencing only after bipolar attachments form (Fig. 1A). How the kinetochore meets this requirement is unclear, because the same protein, Protein phosphatase 1 (PP1), antagonizes both the SAC and error correction. PP1 silences the SAC by dephosphorylating the kinetochore protein KNL1/Spc105 to enable anaphase onset (London et al., 2012; Meadows et al., 2011; Nijenhuis et al., 2014; Rosenberg et al., 2011). It also stabilizes kinetochore-microtubule attachments by dephosphorylating microtubule-binding kinetochore components such as the Ndc80 complex (Liu et al., 2010; Posch et al., 2010). This dual role of PP1 creates the possibility for a harmful cross-talk between SAC silencing and error correction: if PP1 is recruited for SAC silencing before chromosome biorientation, it will inadvertently stabilize syntelic attachments and cause chromosome missegregation. Whether this harmful cross-talk occurs, and how it can be mitigated remains unclear.

**Figure 1.**
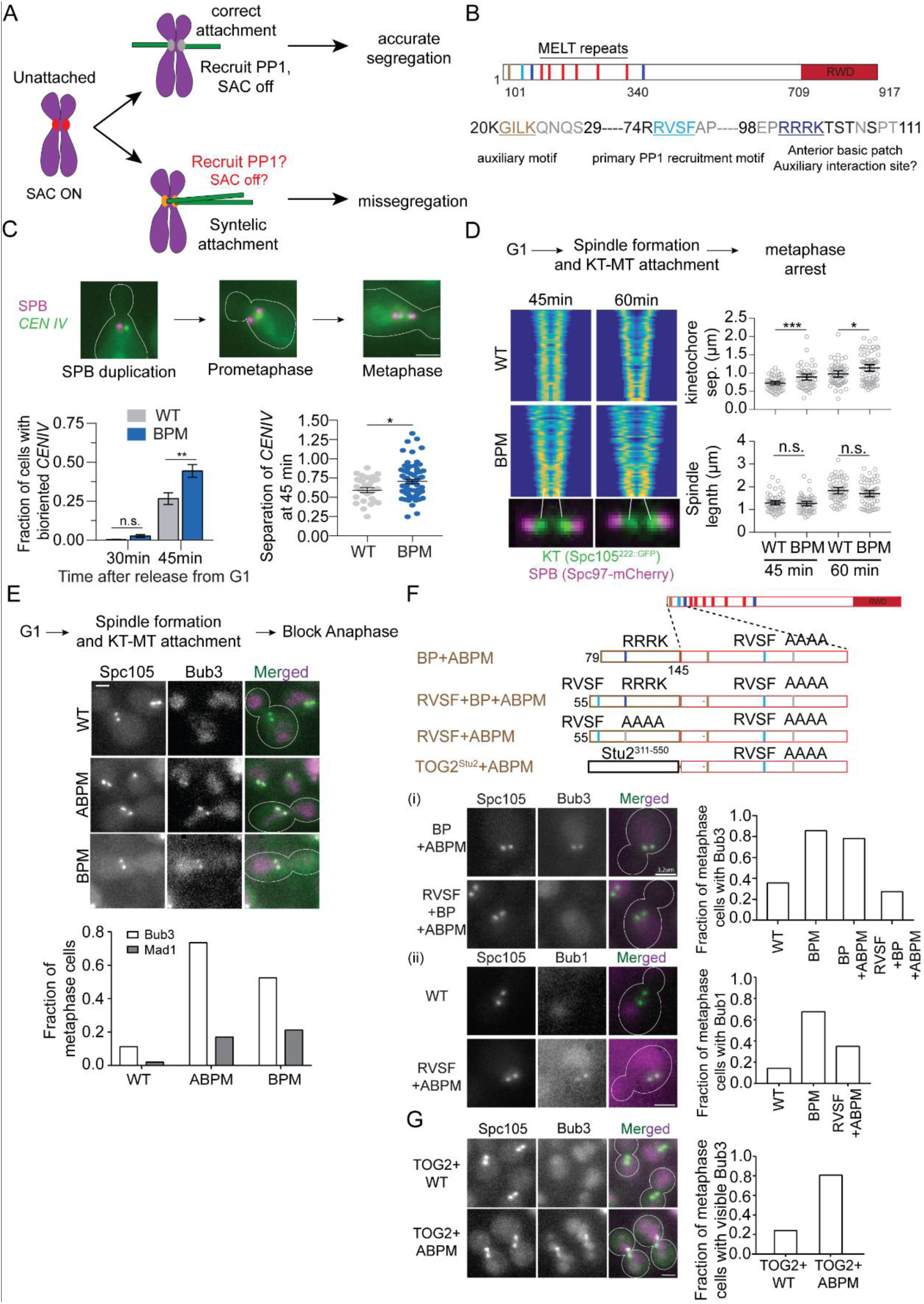
The basic patch near the N-terminus of Spc105 contributes to Glc7 recruitment. (A) Model of how cross-talk between SAC silencing and error correction can interfere with the correction of syntelic attachments and promote chromosome missegregation. **(B)** Functional domains of Spc105 and the amino acid sequence of its N-terminus. **(C)** Representative micrographs of TetO-TetR-GFP spots. Centromere IV (*CEN IV)* achieves biorientation faster in cells expressing Spc105^BPM^ as compared wild-type cells (p ∼ 0.0041 at 45 min. using the Two-way ANOVA test). Sister centromere separation is higher in cells expressing Spc105BPM as compared to wild-type cells even though the spindle length is not (Scale bar∼3.2μm). **(D)** Left panel: V-plots display normalized distribution of kinetochores along the spindle axis for the indicated strains (n > 50 for each time point). Each row of pixels in the plot represents the symmetrized distribution of Spc105^222∷GFP^ or Spc105^BPM,222∷GFP^ along the spindle axis in one cell. Rows are ranked according to spindle length [see methods and (Marco et al., 2013)]. Right top: Average sister kinetochore separation (p=0.0005 and 0.0121 for 45 and 60 min respectively using the unpaired t-test). Right bottom: distance between two spindle poles remains unchanged (p=0.6523 and 0.1932 for 45 and 60 min respectively using the unpaired t-test, from 2 experiments). **(E)** Top: work flow: Middle: representative micrographs of yeast cells expressing the indicated proteins. Scale bar∼3.2μm. Bottom: Frequency of metaphase cells with visible Bub3 and Mad1 at the kinetochores (n = 204, 196 and 179 respectively pooled from 2 experiments for Bub3-mCherry and n= 101, 94 and 123 for Mad1-mCherry). In this and subsequent assays yielding two-category (presence or absence of visible recruitment) scoring data for wild-type and mutant Spc105, we used Fisher’s exact test for the fractions calculated from the total number of observations. p<0.0001 for Bub3-mCherry and < 0.0003 For Mad1-mCherry recruitment. **(F)** Fusion of an extra basic patch with or without RVSF and their effect on Bub3-Bub1 localization in metaphase cells. (i) Left: Representative micrographs (Scale bar∼3.2μm). Right: Bar graph shows the fraction of cells with visible Bub3-mCherry recruitment (n = 90, 70, 96 and 114 for respectively pooled from 2 experiments; *p* < 0.0001 using Fisher’s exact test). (ii) Left: Representative micrographs of cells expressing the indicated proteins (Scale bar∼3.2μm). Right panel: Bar graph shows the fraction of cells with visible Bub1-mCherry recruitment (n = 42, 120 and 103 for respectively pooled from 2 experiments; *p* < 0.0152 using Fisher’s exact test). (G) The effect of TOG2-Spc105 fusions on Bub3-mCherry recruitment to bioriented kinetochores. Left: Representative micrographs (Scale bar∼3.2μm). Right: Bar graph showing fraction of metaphase cells with visible Bub3-mCherry recruitment (n=112 and 144 for TOG2-Spc105 and TOG2-Spc105^ABPM^ respectively, pooled from 2 experiments).

To study the coordination between SAC silencing and error correction, we investigated the importance of regulated PP1 recruitment by Spc105. PP1 recruitment by Spc105 is necessary for SAC silencing, and it is also thought to stabilize kinetochore-microtubule attachments (Hendrickx et al., 2009; Liu et al., 2010; London et al., 2012; Nijenhuis et al., 2014; Rosenberg et al., 2011). Interestingly, Aurora B, the kinase responsible for error correction, downregulates the interaction between Spc105 and PP1. In principle, this regulation can minimize the cross-talk discussed above, but the role of Aurora B is thought to be important mainly for robust SAC in unattached kinetochores (Liu et al., 2010; Nijenhuis et al., 2014). Using cell biological and genetic experiments, we show that the regulation of PP1 recruitment by Spc105 is also critical for chromosome biorientation and accurate chromosome segregation. If PP1 is recruited by Spc105 prematurely, i.e. before sister kinetochore biorientation, it inadvertently stabilizes syntelic attachments and interferes with error correction. We find that a patch of basic residues in the N-terminus of Spc105 makes a regulatable contribution to PP1 recruitment. We also show that phosphoregulation of the activity of the basic patch can improve the accuracy of chromosome segregation. Ultimately, our results demonstrate the presence of a harmful cross-talk between SAC silencing and error correction, explains how it is mitigated in budding yeast, and also uncovers a novel role for the phosphoregulation of PP1 recruitment via Spc105.

## Results and Discussion

### The loss of a conserved basic patch near the N-terminus of Spc105 leads to defective SAC silencing and also improves kinetochore biorientation

The N-terminus of Spc105 contains two known PP1 recruitment sites: the RVSF motif (primary binding site) and the ‘G/SILK’ motif (the secondary site, Fig. 1B). The Aurora B kinase, known as Ipl1 in budding yeast, phosphorylates the RVSF motif, but the abrogation of this phosphoregulation does not appear to have detectable effects on chromosome segregation in budding yeast (Rosenberg et al., 2011). The secondary binding site in budding yeast lacks a phosphorylatable residue. This suggests the existence of a third element involved in the regulation of PP1 recruitment. Indeed, mutation of a patch of four basic patch in the N-terminus of the *C. elegans* homolog of Spc105 results in defective SAC silencing raising the possibility that it could regulate PP1 activity directly or indirectly (Espeut et al., 2012). Spc105 contains two basic patches, an anterior one spanning residues 101-104 and a posterior one spanning residues 340-343. Notably, the anterior conserved patch of four basic residues ‘RRRK’ may be subject to phosphoregulation, because the Serine and Threonine residues located immediately downstream from the basic patch are phosphorylated by mitotic and S-phase kinases [pSYT repository of phosphorylated peptides hosted at the Global Proteome Machine Database; original observations from (Kanshin et al., 2017; Smolka et al., 2007)]. Therefore, to understand the activity of the basic patch, we created the basic patch mutant (BPM) of Spc105, Spc105^BPM^, wherein the basic residues in both basic patches are replaced with non-polar Alanine residues. Mutation of either one or both basic patches showed no adverse effects on cell growth or viability (Fig. S1A). This mutation also did not affect the number of Spc105 molecules in the kinetochore (Fig. S1B).

The earlier study of the basic patch in CeKNL1 in *C. elegans* suggested that it binds to microtubules (Espeut et al., 2012). Therefore, we first tested the basic patches of Spc105 contribute to microtubule binding. Single molecules of recombinant Spc105 phosphodomain (residues 2-455 as a part of 6xHIS-MBP-Spc105^2-455, 222::GFP^) did not detectably interact with Taxol-stabilized porcine microtubules based on Total Internal Reflection Fluorescence microscopy observations (data not shown). Therefore, we tested the binding of Spc105 and Spc105^BPM^ coated microspheres to microtubules. Wild-type 6xHIS-MBP-Spc105^2-455, 222::GFP^-coated microspheres readily bound to microtubules, and whereas 6xHIS-MBP-Spc105^2-455,BPM,222::GFP^-coated microspheres did not bind (Fig. S1B). These results indicate the basic patches of Spc105 can interact with microtubules. However, since single molecules of the phosphodomain do not interact with the microtubule, the basic patches likely contribute only a weak microtubule interaction.

We next analyzed the effect of Spc105^BPM^ on the kinetics of kinetochore biorientation and force generation *in vivo*. Using a centromere-proximal TetO array to visualize centromere IV (*CEN IV*), we quantified the fraction of cells that achieve biorientation 30 and 45 minutes after release from a G1 arrest. After 45 minutes, *CEN IV* achieved biorientation in a significantly larger fraction of cells expressing Spc105^BPM^ compared to wild-type cells (Fig. 1C). Interestingly, the separation between the bioriented *CEN IV* was also significantly higher in cells expressing Spc105^BPM^ (Fig. 1C). We confirmed these results by examining the kinetics of biorientation of all 16 pairs of sister kinetochores. In this experiment, we used with GFP-tagged Spc105 to visualize all kinetochores and quantified their distribution over the spindle using a previously described method (Marco et al., 2013). Cells expressing Spc105^BPM^ bioriented their kinetochores faster as compared to cells expressing wild-type Spc105 (compare the V-plots in Fig. 1D (left) which displays kinetochore distribution of > 50 spindles imaged at the indicated times after release from the G1 arrest). These results show that Spc105^BPM^ speeds kinetochore biorientation, The average distance between the centroids of sister kinetochore clusters was again higher in cells expressing Spc105^BPM^ even though the spindle length was the same (scatterplots in Fig. 1D). This increased separation could indicate higher force generation by the kinetochore. Neither phenotypes can be easily explained by a reduction in microtubule binding affinity. They suggest that the basic patches contribute to a function other than force generation. The basic patch does not affect kinetochore force generation in *C. elegans* embryos and human cells (Espeut et al., 2012; Zhang et al., 2014).

We next assessed whether Spc105^BPM^ impairs SAC silencing, similar the reported function of the basic patch in C. elegans. To do this, we quantified the recruitment of Bub3- or Bub1-mCherry to bioriented kinetochores in metaphase-arrested cells (using *CDC20* repression, see Methods). Bub3 is a key reporter of active SAC signaling. It only binds to phosphorylated MELT motifs in Spc105 as a part of the Bub3-Bub1 complex (Aravamudhan et al., 2015; Hiruma et al., 2015; Ji et al., 2015; London et al., 2012; Primorac et al., 2013). Once stable kinetochore-microtubule attachments form, PP1, Glc7 in yeast, dephosphorylates the MELT motifs to suppress Bub3 recruitment (London et al., 2012). Therefore, in wild-type cells only a small minority of metaphase-arrested yeast cells show detectable Bub3 recruitment at bioriented kinetochores (Fig. 1E). In contrast, under the same conditions in yeast cells expressing either Spc105^BPM^ or Spc105^ABPM^ (wherein only the anterior basic patch is mutated) Bub3-mCherry was on the majority of bioriented kinetochores (Fig. 1E). We next monitored the recruitment of Mad1 to kinetochores under the same conditions. Mad1 binding to the kinetochore is required to activate the SAC. In contrast to Bub3, visible Mad1 localization was seen in a small fraction of the cells expressing either Spc105^BPM^ or Spc105^ABPM^ (Fig. 1E, micrographs and quantification shown in Fig. S1D). Consistent with this observation, the growth rate of Spc105^BPM^ cultures is very similar to the growth rate of wild-type cells indicating little or no delay in cell cycle progression (Fig. S1F).

Given our current understanding of the kinetochore, the abnormal Bub3-mCherry recruitment to kinetochores in metaphase-arrested cells can be explained by two different mechanisms. First, if the basic patch acts solely in microtubule binding, the loss of microtubule binding by Spc105^BPM^ may act like an unattached kinetochore and release the Spc105 from the microtubule lattice thereby bringing the MELT motifs into proximity of the Mps1 kinase which phosphorylates them (Aravamudhan et al., 2015). Alternatively, Spc105^BPM^ may act by a novel mechanism that reduces Glc7 recruitment or activity, and thus suppresses the dephosphorylation of MELT motifs and hence increases Bub3 recruitment. To understand which of these two mechanisms is responsible, we first tested whether the abnormal Bub3 recruitment is suppressed by the re-introduction the basic patch into Spc105^BPM^ at a different location in the N-terminus of Spc105. To do this, we fused a fragment of Spc105 containing the basic patch and its surrounding residues to the N-terminus of Spc105^BPM^ [residues 79-145 (BP+BPM), Fig. 1F i]. This did not suppress the abnormal Bub3 recruitment. As an alternative approach to test the role of microtubule binding in the suppression of Bub3 recruitment, we fused the TOG2 domain (Stu2^311-550^) from Stu2/XMAP215, a known microtubule-binding domain, to the N-terminus of Spc105^BPM^ (Ayaz et al., 2012). Bioriented kinetochores in the majority of the cells expressing TOG2-Spc105^BPM^, but not TOG2-Spc105 recruited Bub3-mCherry suggesting that microtubule binding is not sufficient to suppress the Bub3 recruitment/retention (Fig. 1G). These results suggest that the basic patch may be acting via a novel mechanism to suppress Bub3 recruitment/retention.

The alternative mechanism by which Spc105^BPM^ results in aberrant Bub3-mCherry recruitment is that the basic patch may contribute to Glc7 recruitment. To test this mechanism, we next appended a fragment of Spc105 (residues 55-145) that contained the RVSF motif either with (RVSF+BP+ABPM) or without the basic patch (RVSF+ABPM) to the N-terminus of Spc105^ABPM^ (see schematics at the top of Fig. 1F). The abnormal recruitment of Bub3-Bub1 was suppressed only when the extra RVSF motif was introduced along with the downstream basic patch (RVSF+BP+BPM, Fig. 1F i and ii). These data strongly suggest that the RVSF motif and the basic patch act together to suppress Bub3 recruitment likely through recruiting or activating Glc7 at the yeast kinetochore.

### The basic patch promotes Glc7 recruitment to the kinetochore

To directly test whether the basic patch contributes to Glc7 binding to Spc105 *in vivo*, we used the kinetochore particle pull-down assay to assess the interaction of Glc7 with kinetochores in cells expressing Spc105^BPM^ (Gupta et al., 2018). Briefly, we immunoprecipitated Dsn1-Flag to pull down kinetochore particles from yeast cells expressing either wild-type Spc105^222::mCherry^ or Spc105^BPM,222::mCherry^, and quantified the amount of Spc105 and Glc7-3xGFP co-precipitating with the kinetochore particles (Methods). After normalizing to the levels of Spc105^222::mCherry^ in the co-precipitate, we found that the level of Glc7 associating with kinetochore particles containing Spc105^BPM, 222::mCherry^ was reduced by ∼ 62-85% (two independent experiments, shown in Fig. 2A and Fig. S1E). Thus, the basic patch mutation significantly reduces the interaction of Glc7 with yeast kinetochores, implying that the basic patch cooperates with the RVSF motif to achieve normal levels of Glc7 recruitment in yeast kinetochores.

**Figure 2.**
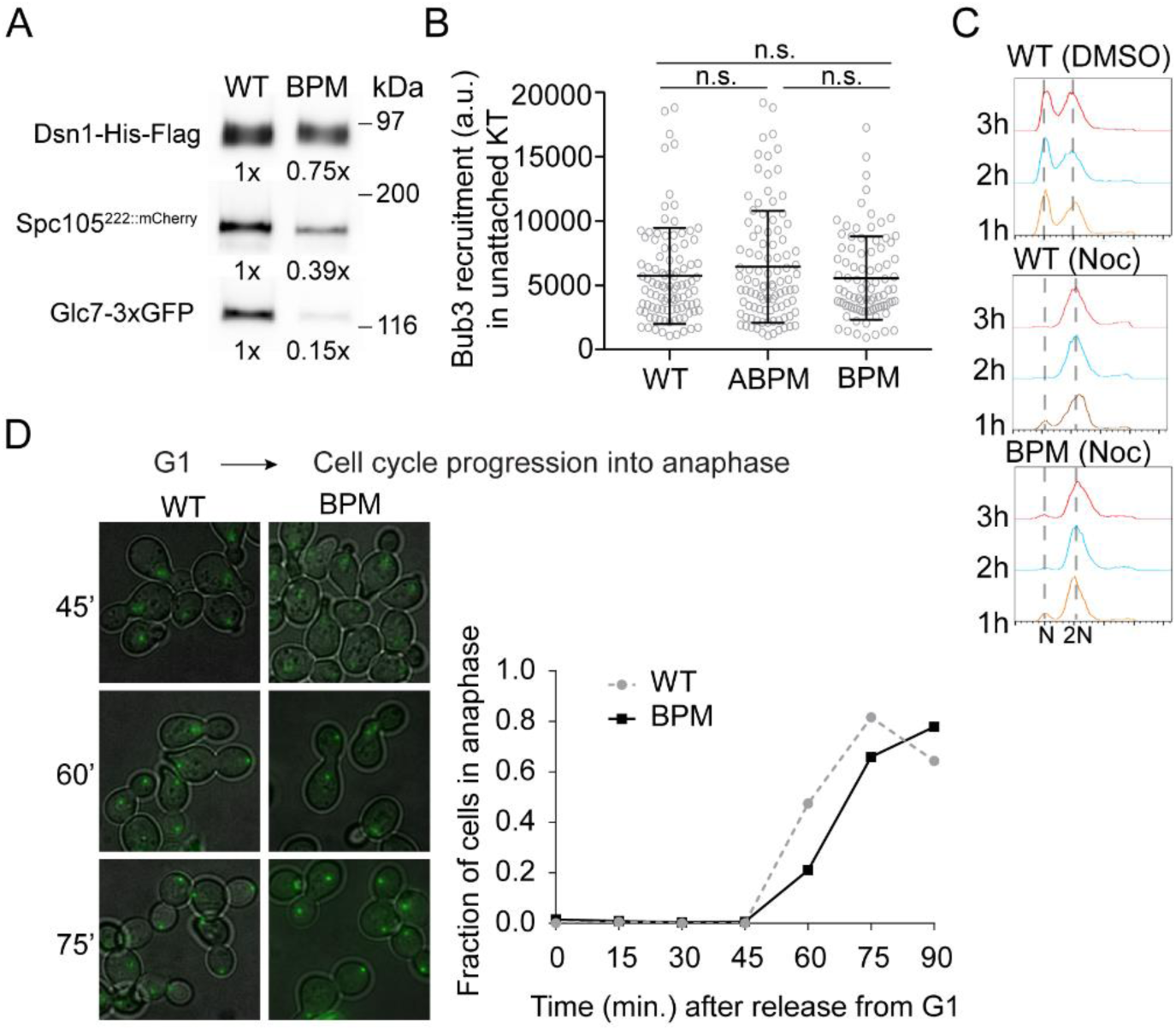
Spc105^BPM^ does not enhance SAC signaling. **(A)** I*n vivo* recruitment of Glc7-3xGFP by kinetochore particles containing either wild-type Spc105 (WT) or Spc105^BPM^ (BPM). Numbers below each lane indicate the band intensity relative to the band intensity for the wild-type Spc105. The reduced co-precipitation of Spc105^BPM^ reflects experimental variation, and not a reduction in the number of kinetochore-bound molecules (see Fig. S1B). **(B)** Bub3-mCherry recruitment by unattached kinetochore clusters in strains expressing the indicated Spc105 variant (mean ± s.d.; p=0.2501 obtained from one-way ANOVA test, n = 97, 97, and 88 respectively pooled from 2 experiments). **(C)** Flow cytometry of the DNA content per from cultures treated with Nocodazole. The 1n and 2n peaks correspond to G1 and G2/M cell populations respectively (representative histograms from 2 trials). **(D)** Left: representative transmitted light and fluorescence micrographs of yeast cells at indicated time points after release from a G1 arrest. The chart the fraction of cells entering anaphase at the indicated time after release from the G1 arrest (data from pooled from two experiments). It should be noted that the drop in the fraction of anaphase cells in the wild-type culture at the last time point is because many cells enter the next cell cycle. Scale bar∼2.0μm.

### Spc105^BPM^ does not enhance SAC signaling

The lack of Mad1 recruitment to bioriented kinetochores in cells expressing Spc105^BPM^ and normal growth rate indicated that the defect in SAC silencing does not translate into a cell cycle delay (Fig. 1E and S1D). However, because of the involvement of the basic patch in Glc7 recruitment, we analyzed in detail whether Spc105^BPM^ affects SAC signaling and its silencing. We treated cells expressing either wild-type Spc105 or Spc105^BPM^ with the microtubule poison nocodazole to depolymerize the spindle and quantified the amount of Bub3-mCherry recruited by unattached kinetochores. In both cases, unattached kinetochores recruited similar amounts of Bub3 indicating that SAC signaling was unaffected by the mutation (Fig. 2B). This is consistent with our prior finding that Glc7 activity has little influence on Bub3 recruitment in unattached kinetochores (Aravamudhan et al., 2016). We next used flow cytometry to quantify changes in the DNA content of cells treated with nocodazole over time. Spc105^BPM^ had no detectable effect on the DNA content, indicating that the strength of the SAC was not detectably affected by the mutation (Fig. 2C). Finally, we tested whether Spc105^BPM^ delays the metaphase to anaphase transition by monitoring its timing in a synchronized cell population. Anaphase onset was marginally delayed in cells expressing Spc105^BPM^, similar to the effect of a similar mutation in *C. elegans* (Fig. 2D). Together, these results show that Spc105^BPM^ has minimal impact on SAC signaling and causes at most a minor delay in SAC silencing.

### Spc105^BPM^ significantly improves chromosome segregation in a Sgo1-independent manner

According to the current understanding, reduced Glc7 recruitment by Spc105^BPM^ should reduce the dephosphorylation of microtubule-binding kinetochore proteins, and therefore destabilize kinetochore-microtubule attachments. This should in turn delay kinetochore biorientation and reduce sister kinetochore separation. The faster kinetics of kinetochore biorientation in cells expressing Spc105^BPM^ (Fig. 1C) is inconsistent with this model. One potential explanation for the observed phenotype is that the higher Bub1 level at the kinetochore promotes Shugoshin (Sgo1) recruitment (Fig. S2A) and enhances sister centromere cohesion and Aurora B activity, and thus promotes sister kinetochore biorientation (Kawashima et al., 2010a; Kawashima et al., 2010b; Peplowska et al., 2014; Salic et al., 2004). To test this possibility, we examined interactions between the genes involved in the Sgo1-mediated error correction pathway and Spc105^BPM^ using the benomyl sensitivity assay (see Fig. 3 top panel for a schematic of the error correction pathway). In this assay, yeast cells are grown on media containing low doses of the drug benomyl. In the presence of benomyl, microtubule dynamicity increases, and as a result, the spindle becomes short, oscillatory movements of bioriented kinetochores dampen, and centromeric tension is significantly reduced (Pearson et al., 2003). To proliferate under these conditions, yeast cells must possess robust SAC signaling and error correction mechanism. Mutant strains that are defective in either SAC signaling (e.g. *mad2Δ*) or error correction (e.g. *sgo1Δ*) grow poorly or are inviable specifically on benomyl-containing media (Fig. 3).

**Figure 3.**
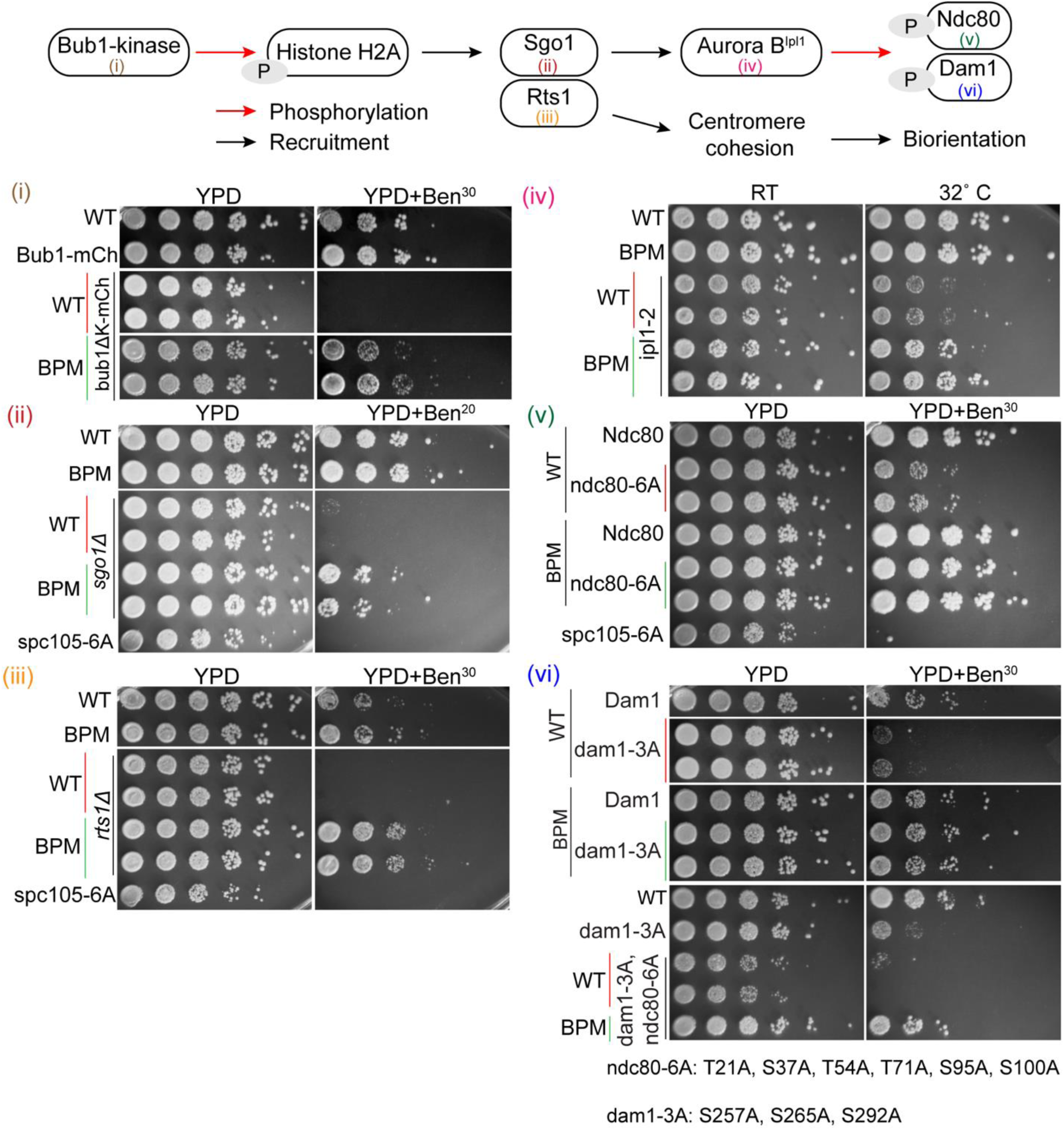
Spc105^BPM^ improves the error correction pathway independently of Sgo1. Top: Schematic of the sister centromere cohesion and biorientation pathway in budding yeast. Bottom: Suppression of benomyl sensitivity by Spc105 mutations, Serial dilutions of yeast cells spotted on either rich media or media containing either 20 or 30 μg/ml Benomyl. WT at the top of each plating indicates wild-type strain included as a positive control. WT, BPM, or ABPM in other rows refers to the Spc105 allele. *spc105-6A*, wherein all six MELT motifs are rendered non-phosphorylatable, was used as the negative control.

Yeast strains carrying either the *bub1^Δkinase^* mutation, wherein Bub1 lacks its kinase domain, or *sgo1Δ*, are impaired in error correction, but competent in SAC signaling (Fig. S2B). Because of the error correction defect, they can grow on normal media, but they cannot grow on benomyl-containing media (Fig. 3 i-ii). To test the role of Glc7 binding in error correction, we examined the genetic interactions of the Spc105 mutants and chimeras with *bub1^Δkinase^* or *sgo1Δ.* Strikingly, *spc105^BPM^* displayed a positive genetic interaction with both these strains: the double mutants *spc105^BPM^ bub1^Δkinase^* and *spc105^BPM^ sgo1Δ* grew robustly on benomyl-containing media (Fig. 3 i-ii). These results indicate that reduced PP1 recruitment to the kinetochore via Spc105 results in improved chromosome segregation in cells with an impaired error correction mechanism. Using the positive genetic interaction between *spc105^BPM^* and *sgo1Δ* as a functional readout, we confirmed that the mutation of the anterior basic patch (Spc105^ABPM^), but not the posteriorly located basic patch (Spc105^PBPM^), suppresses the benomyl-lethality of *sgo1Δ* cells (Fig. S2C). Moreover, consistent with the model that the basic patch works in concert with the RVSF motif, RVSF+APBM, but not RVSF+BP+ABPM, suppressed the benomyl-sensitivity of *bub1^Δkinase^* (Fig. S2D, see Fig. 1F for schematic of Spc105 chimeras). These results indicate recruitment of Glc7 by the RVSF motif and the basic patch working together inhibits error correction.

To further elucidate how Spc105^BPM^ promotes accurate chromosome segregation, we studied genetic interactions of *spc105^BPM^* and genes involved in kinetochore biorientation and error correction (see the pathway schematic in Fig. 3A). *spc105^BPM^* suppressed the benomyl-lethality of the deletion of the gene *RTS1* (Fig. 3 iii), which encodes a regulatory subunit of the Protein Phosphatase 2A involved in sister chromatid cohesion (Peplowska et al., 2014; Xu et al., 2009). *spc105^BPM^* also suppressed the temperature sensitivity of a strain expressing *ipl1-2* (Ipl1-H352Y) that has a weaker kinase activity (Fig. 3 iv). Finally, *spc105^BPM^* suppressed the benomyl sensitivity of strains expressing *ndc80-6A* and *dam1-3A*, alleles of the microtubule-binding proteins Ndc80 and Dam1, wherein all but one of the known Ipl1 phosphorylation sites in the respective are rendered non-phosphorylatable (Fig. 3 v and vi), mutated phosphosites indicated at the bottom, (Cheeseman et al., 2002; Lampson et al., 2004; Pinsky et al., 2006; Tanaka et al., 2002). These mutations normally result in hyper-stable kinetochore-microtubule attachments, which interfere with error correction (Akiyoshi et al., 2009). Strikingly, Spc105^BPM^ suppresses these defects indicating that the accuracy of chromosome segregation is significantly improved by the loss of the basic patch.

These results reveal that *spc105^BPM^* improves chromosome segregation. The suppression of the benomyl-sensitivity of the *rts1Δ* mutant by *spc105^BPM^* is especially noteworthy. *rts1Δ* only affects sister centromere cohesion; it does not affect centromeric recruitment of either Sgo1 or Ipl1 (Peplowska et al., 2014). Yet, reduced recruitment of Glc7 via Spc105 leads to significant improvement in chromosome segregation in the *rts1Δ* mutant. This observation, together with the suppression of the *ndc80-6A dam1-3A* double mutant phenotype by *spc105^BPM^* shows that *spc105^BPM^* reduces necessity of a stringent Ipl1/Aurora B kinase mediated error correction pathway for ensuring accurate chromosome segregation.

### Spc105^BPM^ improves the accuracy of chromosome segregation in *sgo1Δ* cells

The benomyl lethality of yeast strains carrying mutations in centromeric, kinetochore and SAC proteins is due to a lower accuracy of chromosome segregation. Therefore, the suppression of this benomyl lethality by *spc105^BPM^* of these mutants suggests that *spc105^BPM^* improves the accuracy of chromosome segregation. To test this prediction, we monitored chromosome segregation by fluorescently marking *CEN IV* using TetO repeats in *sgo1Δ* cells. We used this mutant strain background, because chromosome missegregation is rare and not easily quantified in wild-type cells under normal growing conditions (Verzijlbergen et al., 2014). We first destroyed the spindle by treating cells with nocodazole, and then monitored the kinetics of *CEN IV* biorientation. Consistent with our earlier observations, TetO spots marking *CEN IV* separated from one another faster in cells expressing Spc105^BPM^ as compared to wild-type cells (Fig. 4A, right). Importantly, the frequency of *CEN IV* missegregation was significantly reduced in the cells expressing Spc105^BPM^, indicating that the efficiency of chromosome segregation was enhanced as compared to *sgo1Δ* cells. Thus, Spc105^BPM^ improves the kinetics of kinetochore biorientation and reduces chromosome segregation errors.

**Figure 4.**
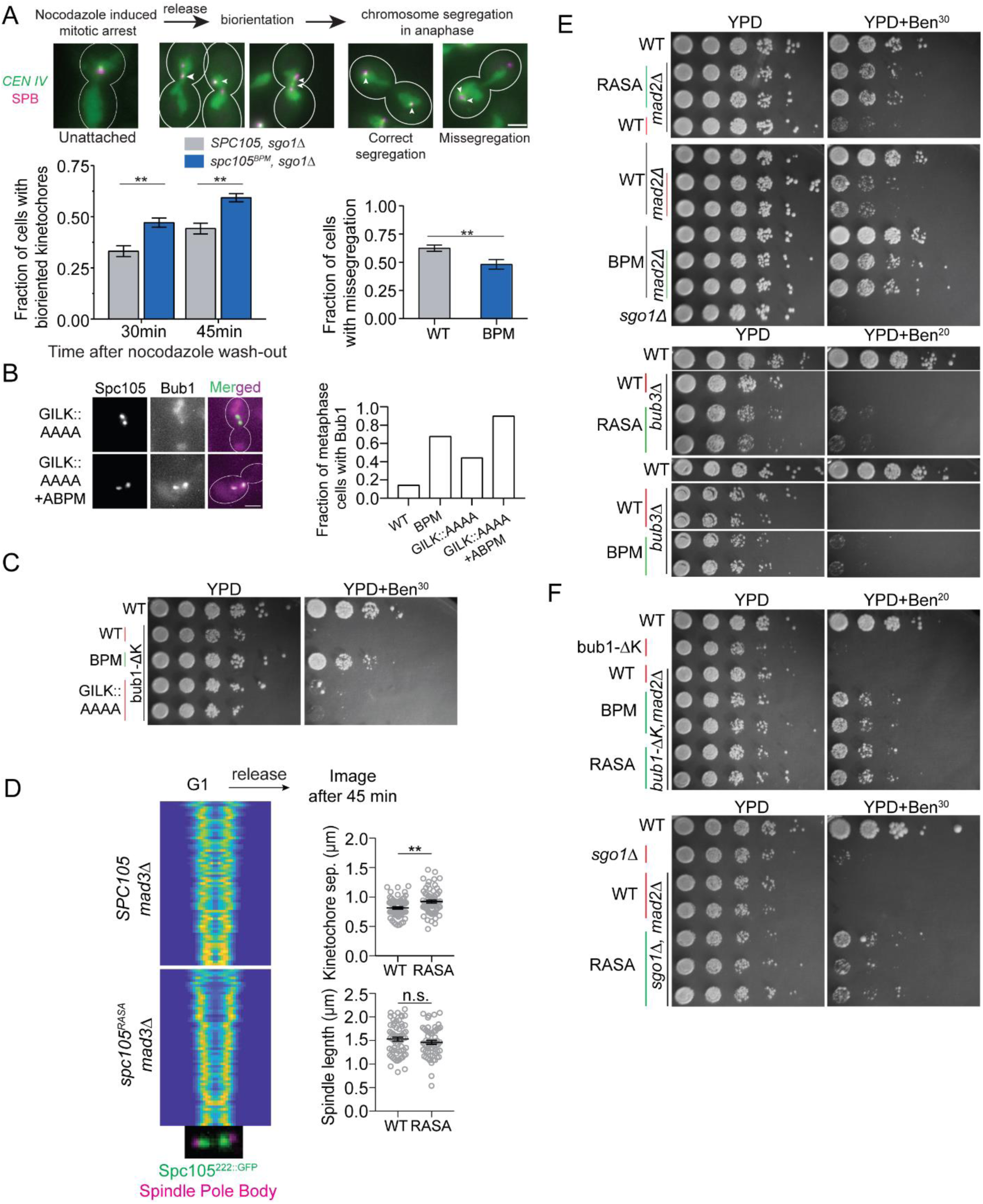
Glc7 recruited by the RVSF motif in Spc105 is not required for error correction. **(A)** Top: workflow used to study chromosome biorientation. Arrowheads show the positions of *CEN IV* foci within the spindle axis in each cell (Scale bar∼3.2μm). Bottom left: Fraction of cells with bioriented *CEN IV* at the indicated time after nocodazole wash-out (Two-way ANOVA test revealed p=0.0066 at 30 min, p=0.0043 at 45 min. n = 434 and 464 at 30 and 45 minutes respectively for Spc105^WT^. n=458 and 411 at 30 and 45 minutes respectively for Spc105^BPM^, accumulated from 3 repeats). Bottom right: Fraction of anaphase cells with chromosome IV mis-segregation quantified 45 minutes post wash-out (p=0.0079 according to unpaired t-test). (**B**) The fraction of metaphase cells with detectable Bub1-mCherry recruitment to bioriented kinetochores in cells expressing the indicated Spc105 variant (p < 0.0015 for all mutant-wild-type comparisons using Fisher’s exact test).(**C**) Spotting assay of the indicated strains on benomyl-containing media. **(D)** Separation between sister kinetochore clusters in the indicated strains and at the indicated time after release from a G1 arrest (analysis performed as in 1C, >48 cells displayed). Right top panel: separation between two sister kinetochores in metaphase (p=0.0014, unpaired t-test). Right bottom panel: distance between two sister SPBs in metaphase (p= 0.2285, obtained from unpaired t-test). **(E and F)** Spotting assay of the indicated strains on benomyl-containing media.

### Glc7 recruited via the RVSF motif in Spc105 is not required for stabilization of kinetochore-microtubule attachment

Using the results so far, we can conclude that: (1) Spc105^BPM^ recruits significantly lower amount of Glc7 to the kinetochore, and (2) cells expressing Spc105^BPM^ exhibit improved accuracy of chromosome segregation. Together, these conclusions imply that a weakened Spc105-Glc7 interaction impairs SAC silencing but improves the chromosome biorientation and the accuracy of chromosome segregation. To test this further, we determined whether the mutation of the canonical Glc7 binding sites in Spc105, the GILK and RVSF motif, results in similar phenotypes as Spc105^BPM^. Similar to Spc105^BPM^, cells expressing Spc105^GILK::AAAA^ showed abnormal recruitment of Bub3-Bub1 to bioriented kinetochores (Fig. 4B). Furthermore, the effect of basic patch and the GILK motif mutations together produced an additive effect: the fraction of metaphase cells recruiting Bub3-Bub1 increased in this double mutant (Fig. 4B). However, *spc105^GILK::AAAA^* by itself did not suppress the benomyl sensitivity of *bub1^Δkinase^* strains (Fig 4C), suggesting that the GILK motif makes a relatively small contribution to Glc7 recruitment as compared to the basic patch.

To understand if Glc7 recruitment via the RVSF motif in Spc105 is required for kinetochore biorientation, we use the *spc105^RASA^* allele which completely blocks Glc7 binding to Spc105. Because this mutation makes Spc105 unable to recruit Glc7, cells arrest in metaphase and cannot grow. Therefore, we used the *spc105^RASA^ mad3Δ* strain to allow growth (Rosenberg et al., 2011). As before, we analyzed the kinetics of kinetochore biorientation by imaging the distribution of fluorescently labeled kinetochores over the spindle as cells progressed from G1 into the cell cycle. Similar to Spc105^BPM^, kinetochore clusters in *spc105^RASA^ mad3Δ* cells were separated by a larger distance as compared to *mad3Δ* cells (Fig. 4D). This simple assay confirms that the recruitment of Glc7 via Spc105 is not required for kinetochore biorientation.

A weakened interaction between Glc7 and PP1 may also improve kinetochore biorientation simply by delaying SAC silencing, and thus providing additional time for kinetochore biorientation (Munoz-Barrera et al., 2015). To test this, we exploited the SAC-deficient *spc105^RASA^* strains and the benomyl sensitivity assay as a functional test of the accuracy of chromosome segregation. Due to the inactive SAC in these strains (Fig. S2E), any improvement in the accuracy of chromosome segregation must occur due to improved sister kinetochore biorientation and error correction. Strikingly, *spc105^RASA^ mad2Δ* grew robustly on benomyl-containing media, even though *mad2Δ* grew poorly under the same condition (Fig. 4E). *spc105^RASA^* also partially suppressed the benomyl lethality of *bub3Δ.* The milder rescue likely reflects the additional defects due to *bub3Δ* that include lower Sgo1 recruitment to the centromere and defects in APC/C function [Fig. 4E (Yang et al., 2015)]. Consistent with these results, *spc105^BPM^* suppressed the benomyl sensitivity due to *mad2Δ*, and mildly suppressed the benomyl lethality due to *bub3Δ*. Most strikingly, the triple mutants *spc105^RASA^ mad2Δ bub1^Δkinase^* and *spc105^RASA^ mad2Δ sgo1Δ* also grew on benomyl-containing media (Fig. 4F). Thus, the improved accuracy of chromosome segregation observed upon the weakening or loss of the Glc7-Spc105 interaction is not because of delayed SAC silencing.

In conclusion, Glc7 recruitment by the RVSF motif of Spc105 is not required for stabilizing kinetochore-microtubule attachments. On the contrary, our results imply that the Glc7 recruited by Spc105 can stabilize syntelic kinetochore-microtubule attachments and interfere with the error correction mechanism. When the RVSF motif is inactivated, these deleterious effects are eliminated, and this enables efficient chromosome biorientation even when Ipl1/Aurora B activity is reduced, or its substrates contain only one phosphorylation site each (Fig. 3).

### Artificial tethering of PP1 to Spc105 demonstrates the necessity of temporal regulation of the PP1-Spc105 interaction for chromosome biorientation

To directly test whether Glc7 recruitment via Spc105 interferes with chromosome biorientation, we used rapamycin-induced dimerization of the Fkbp12 and Frb domains to artificially tether Glc7 at the N-terminus of Spc105 (Haruki et al., 2008). In this experiment, we tethered Glc7 close to its normal binding sites in Spc105 by adding rapamycin to the growth media either before or after sister kinetochore biorientation, and then monitored its effect on kinetochore biorientation and attachment. These cells express both Spc105 and Frb-Spc105 to avoid tethering abnormally high amounts of Glc7.

We first monitored the effect of Glc7 tethering to Spc105 prior to chromosome biorientation. For this, we arrested cells in mitosis by treating them with nocodazole, and then tethered Glc7 to Spc105 by treating the cells with rapamycin (work flow depicted at the top of Fig. 5A). In nocodazole-treated cells, Bub3 recruitment to unattached kinetochores remained unchanged even after rapamycin treatment, indicating that SAC signaling remained unaffected by the treatment (Fig. S3A). We next released these cells from the mitotic block and examined kinetochore distribution over the mitotic spindle 30 minutes after nocodazole wash out. To ensure that the potentially faster SAC silencing by the tethered Glc7 does not induce premature anaphase onset, we also blocked anaphase-onset in these cells by depleting Cdc20. After 30 minutes, the majority of untreated cells displayed normal spindle morphology with two bioriented kinetochore clusters (Fig. 5A, left). In contrast, most rapamycin-treated cells showed abnormal kinetochore-microtubule attachment and spindle morphology (Fig. 5A, bar graph). The morphological defects included: (1) unequal distribution of kinetochores between the two spindle pole bodies (kinetochore asymmetry, Fig. 5A i, quantified by calculating the absolute difference between the normalized intensity of the brightest pixel in each spindle half), (2) unaligned kinetochores along the spindle axis, (3) reduced separation between kinetochore clusters and spindle pole bodies (0.27±0.02 μm instead of the normal 0.43±0.11 μm mean ± s.e.m., Fig. S3B), (4) increased separation between sister kinetochores (Fig. 5A iii) and (5) abnormally long spindles (Fig. 5A iv). The unequal distribution of kinetochores between the two spindle pole bodies is a hallmark of defective chromosome biorientation (Marco et al., 2013). The significant increase in the spindle length also indicates that there is a smaller number of bioriented chromosomes opposing the outward forces generated within the spindle (Bouck and Bloom, 2007).

**Figure 5.**
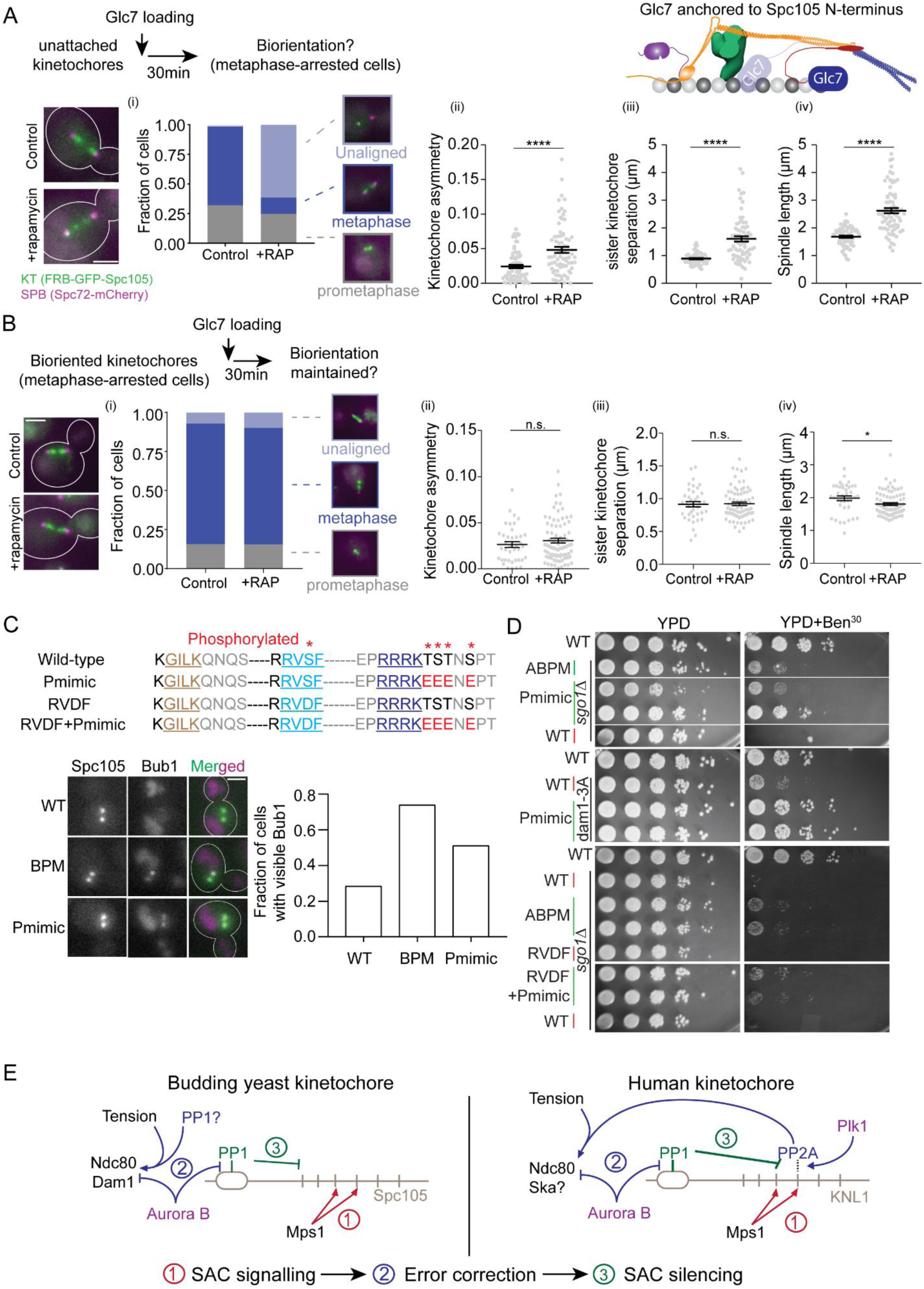
Delayed recruitment of Glc7 for SAC silencing is necessary for the establishment of bipolar chromosome attachment and accurate chromosome segregation. **(A)** Top: the work flow used to tether Glc7 at the N-terminus of Spc105 under prophase-like conditions. Cartoon at the right: 1-D schematic of the yeast kinetochore. Bottom left: micrographs depicting the indicated proteins. Scale bar ∼ 3.2μm. Bottom right: (i) Bar graph depicts the scoring of cells based on spindle morphology (key provided on the right, n = 126 and 89 for control and rapamycin treated samples respectively, pooled from three technical repeats performed on two biological replicates). (ii-iv) Scatter plots display the indicated quantity in untreated control and rapamycin-treated cells. The data is presented as mean ± s.e.m. (n = 88 and 91 control and rapamycin treated cells respectively, data were pooled from three experimental repeats. p<0.0001 with unpaired t-test). (iii) (n = 87 control and rapamycin treated cells pooled from three experimental repeats, p<0.0001 using unpaired t-test) (iv) (n = 127 and 153 control and rapamycin treated cells, pooled from three experimental repeats; p<0.0001 according to unpaired t-test). **(B)** The work-flow for tethering Glc7 to the N-terminus of Spc105 in metaphase-arrested cells. Bottom left: micrographs depicting the indicated proteins. Scale bar∼3.2μm. Bottom right: (i) Bar graph indicates the scoring of cells based on spindle morphology (key provided on the right; n = 70 and 142 for control and rapamycin treated cells respectively pooled from three technical repeats). (ii-iv) Scatter plots display the indicated quantity in untreated control and rapamycin-treated cells. The data is shown as mean± s.e.m. In (ii), 39 and 80 control and rapamycin treated cells pooled from two experimental repeats; p=0.3298, acquired from unpaired t-test. In (iii) and (iv), 38 and 80 cells control and rapamycin treated cells, data pooled from two experimental repeats. In (iii) p=0.8844, obtained from unpaired t-test. In (iv) p=0.0244 from the unpaired t-test showed **(C)** Top: Residues downstream from the basic patch that are known to be phosphorylated are marked with an asterisk at the top. Lower sequences display the mutants used in this study. Bottom: Representative images and quantification of the fraction of metaphase cells with detectable Bub1-mCherry recruitment to bioriented kinetochores (p-value using Fisher’s exact test is <0.0005 for the comparisons displayed on the graph). **(D)** Spotting assay of the indicated Spc105 mutants on benomyl-containing media. **(E)** Left: Model mechanisms that mitigate harmful cross-talk between the SAC silencing and error correction in budding yeast kinetochore. Right: Schematic of the proposed model for the human kinetochore. Dashed line indicates indirect recruitment of PP2A via Mad3/BubR1, which is promoted by the Plk1 kinase.

We next studied the effects of Glc7 tethering to Spc105 after the process of chromosome biorientation is complete (Fig. 5B, schematic at the top). For this, we first repressed *CDC20* expression to block anaphase onset providing sister kinetochores sufficient time to form bipolar attachments, and then added rapamycin to tether Glc7 to the N-terminus of Spc105. In this case, Glc7 tethering had no discernible effects on the metaphase spindle (scatterplots in Fig. 5B): the spindle morphology was nearly identical with or without rapamycin-treatment, kinetochores were symmetrically distributed in two distinct clusters, and spindle length decreased slightly. Thus, Glc7 tethering to Spc105 after kinetochore biorientation does not adversely affect kinetochore-microtubule attachments.

It should be noted that a prior study has shown that Glc7 fusion to the N-terminus of Spc105^RASA^ generates a viable strain with intact SAC signaling, which is inconsistent with the results above (Rosenberg et al., 2011). However, we found that chromosome biorientation was significantly delayed in these cells compared to a wild-type strain of the same strain background, likely because a significant number of cells showed unaligned and unattached kinetochores (Fig. S3C). This strain was also benomyl-sensitive. Therefore, the strain expressing the Glc7-Spc105^RASA^ fusion suffers from defects in chromosome alignment and segregation.

These experiments demonstrate that premature recruitment of Glc7, prior to kinetochore biorientation, strongly interferes with kinetochore biorientation by stabilizing syntelic attachments. However, Glc7 recruitment after kinetochore biorientation does not detectably affect kinetochore-microtubule attachment. These data strongly suggest that the recruitment of Glc7 via Spc105 must be delayed until sister kinetochores establish bipolar attachments.

### Phosphoregulation can mitigate the cross-talk between SAC silencing and error correction

As mentioned previously, the Serine and Threonine residues immediately downstream from the basic patch are phosphorylated by S-phase and mitotic kinases (pSYT accessed at http://gpmdb.thegpm.org/psyt/index.html, also see Methods). This phosphorylation can interfere with the activity of the basic patch to reduce the recruitment of Glc7 via Spc105. Therefore, to test whether the phosphorylation phenocopies Spc105^BPM^, we replaced the reported phosphorylated residues (marked with an asterisk in Fig. 5C) with negatively charged residues, expecting that this will neutralize the electrostatic charge of the basic patch (Fig. 5C, Top panel). Similar to Spc105^BPM^, Bub3-mCherry recruitment to bioriented kinetochore clusters was elevated in cells expressing the phosphomimic allele (Fig 5C, bottom panel). This allele suppressed the benomyl sensitivity of *sgo1Δ* as well as the non-phosphorylatable *dam1-3A* allele, again suggesting a reduced reliance on the error correction mechanisms for sister kinetochore biorientation (Fig. 5D). Importantly, a phosphomimic mutation in the RVSF motif alone did not rescue the benomyl sensitivity of *sgo1Δ* cells (Fig. 5D, lower panel). These data reveal that the phosphorylation of residues downstream from the basic patch by mitotic kinases can reduce Glc7 recruitment to the kinetochore and allow effective kinetochore biorientation.

In summary, our results uncover a harmful cross-talk that can occur between SAC silencing, and chromosome biorientation and error correction. This cross-talk exists, because the set of core kinetochore components involved in these functions are located in close proximity, and because the two functions are regulated by PP1 (Aravamudhan et al., 2014; Aravamudhan et al., 2015). If the Glc7 required for SAC silencing is recruited by Spc105 prematurely – prior to chromosome biorientation, it inadvertently stabilizes syntelic attachments. The experiments involving Spc105^BPM^ and Spc105^RASA^ demonstrate that the weakening of Glc7 recruitment via Spc105 prior to metaphase mitigates this cross-talk. Weakened Spc105-Glc7 interaction allows the error correction mechanism to resolve syntelic attachments, enabling sister kinetochores to achieve bipolar attachments more quickly. Ultimately, these two effects result in accurate chromosome segregation. Delayed SAC silencing resulting from the weakened Spc105-Glc7 interaction plays little or no role in the improved chromosome segregation as demonstrated by the ability of *spc105^RASA^* to suppress the benomyl sensitivity of *mad2Δ* and *bub3Δ* mutants.

We propose that the phosphoregulation of the Spc105-Glc7 interaction minimizes the harmful cross-talk between SAC silencing and error correction. The experiments involving the phosphomimic allele of Spc105, which phenocopies the positive genetic interactions between *spc105^BPM^* and *sgo1Δ* and *dam1-3A*, show that phosphoregulation can establish the required temporal regulation of the Glc7-Spc105 interaction. Efficient kinetochore biorientation in *spc105^RASA^* cells also show that the Glc7 activity required for attachment stabilization is derived from a Spc105-independent source. This Glc7 activity may come from diffusive interactions between Glc7 and the kinetochore. A recent study suggests that the motor protein Cin8 transports Glc7 to the kinetochore (Suzuki et al., 2018). However, it is unclear how this mechanism can deliver Glc7 preferentially to kinetochores with correct, but not incorrect, attachments. A dedicated Glc7 recruitment mechanism may not even be necessary. The inter-centromeric tension generated by sister kinetochores will also selectively stabilize bipolar attachments (Akiyoshi et al., 2010; Franck et al., 2007).

Similar mechanisms can be expected to bring about the ordered execution of chromosome biorientation first and then SAC silencing to ensure accurate chromosome segregation in human cells. Even though a more complex network of three kinases regulates these two processes (Fig. 5E, right panel), their targets including KNL1 are similar (Etemad et al., 2015; Hiruma et al., 2015; London et al., 2012; Nijenhuis et al., 2014; Suijkerbuijk et al., 2012). Aurora B regulates PP1 recruitment by phosphorylating the RVSF motif and the SILK motif, which interestingly, lies directly adjacent to the basic patch in KNL1 (Liu et al., 2010; Welburn et al., 2010). A recent *in vitro* study found that Knl1 binding to the microtubule and PP1 is mutually exclusive, and proposed a model wherein microtubule-binding contributes to regulated PP1 binding to KNL1 (Bajaj et al., 2018). However, this model does not explain why microtubule-binding by KNL1 is needed in the first place if: (a) PP1 binding is inhibited by phosphorylation in unattached kinetochores, where microtubules are absent, and (b) the stronger binding PP1 is expected to displace the microtubule from KNL1 anyway, even when microtubules are present. The true functional significance of the microtubule-binding activity of basic patch in KNL1, which does not contribute to kinetochore force generation, needs to be fully understood (Espeut et al., 2012; Zhang et al., 2014). Our findings suggest a new model (Fig. 5E, right panel), wherein Aurora B suppresses PP1 recruitment by KNL1 to ensure that error correction is completed before PP1-mediated stabilization of bipolar attachments and SAC silencing takes place. In line with this, PP1 recruitment peaks in the human kinetochore only in metaphase, after kinetochores have established bipolar attachments (Liu et al., 2010).

## Acknowledgements

This work was funded by the RO1-GM-088908 to APJ. We would like to thank Mara Duncan for sharing reagents. We thank Rena Evans and Sue Biggins for guidance and training for performing yeast kinetochore particle pull-down experiments, and for providing key reagents.

We thank members of the Joglekar lab and the Duncan lab for their support and constructive comments.

## Methods

### Plasmid and Strain construction

Budding yeast strains and plasmids utilized in this study are listed in Supplementary Tables S1 and S2 respectively. Strains containing multiple genetic modifications were constructed using standard yeast genetics. GFP(S65T) and mCherry tagged to proteins were used to localize kinetochores by microscopy. The C-terminal tags and gene deletion cassettes were introduced at the endogenous locus through homologous recombination of PCR amplicons (Jürg Bähler et al., 1998). A 7-amino-acid linker (sequence: ‘RIPGLIN’) separates the tag (GFP, mCherry) from the C-terminus of the tagged-protein. We have previously observed that the intensity of mCherry-tagged kinetochore proteins varies significantly from strain to strain due to inherent variability of the brightness of mCherry. Therefore, we created all Bub3-mCherry and Mad1-mCherry strains by crossing a specific transformant of the Bub3-mCherry or Mad1-mCherry with strains expressing wild-type Spc105 or its mutants. Mad1-mCherry is always accompanied with *nup60Δ* to prevent it from localizing to the nuclear envelopes (Scott et al., 2005).

Every Spc105 chimera contains a 397 bp upstream and a 250 bp downstream sequence as promoter (*prSPC105*) and terminator (*trSPC105*) respectively. We inserted 711 bp GFP (S65T) fragment (237 amino acids) at 222^nd^ amino acid position of Spc105 by sub-cloning by introducing an extra *Bam*HI site (Gly-Ser) at the upstream and *Nhe*I site (Ala-Ser) at the downstream of the GFP fragment. To insert mCherry using the same *Bam*HI and *Nhe*I sites, we used a 705 bp mCherry fragment (235 amino acids) which was codon-optimized for yeast. We introduced mutations in basic patches (amino acid 101-104 RRRK, 340-343 KRRK) and phosphorylation sites (S77, amino acid 105-107 and S109) by subcloning as well. We incorporated anterior basic patch mutation by initially incorporating an extra *Bsp*EI site that was later removed by site directed mutagenesis to create pAJ525. To create posterior basic patch mutation, we introduced a silent mutation that create *Mlu*I site, which aided us to introduce the mutation at amino acid 340-343. Likewise, we introduced a silent mutation creating *Sac*II site 9 bp upstream of phosphorylation sites (amino acid 105-107 TST, S109) which helped to build the phosphomimetic chimeras. To construct Spc105 chimeras with N-termini fusion (Spc105 79-145, Spc105 55-145 and TOG2), we introduced an extra *Bsp*EI site (Ser-Gly) at the upstream and *Mlu*I site (Thr-Arg) at the downstream, so that we can integrate the fragments after the start codon of Spc105. TOG2 fragment also contains a C-terminal linker peptide of AGGA. FRB-GFP fusion in plasmid pAJ603 consists of 279 bp FRB (93 amino acids) and 711 bp GFP which are linked with each other by a 30 bp fragment which codes for LESSGSGSGS. The whole fragment is ligated within *Bam*HI-*Nhe*I sites after the start codon of Spc105.

To build haploid strains expressing Spc105^222::GFP^ alleles (wild-type or mutant with GFP inserted at amino acid 222), first, we deleted a wild-type genomic copy of *SPC105* in a diploid strain of YEF473 to form the strain AJY3278 *(SPC105/spc105Δ::NAT1)*. Then we transformed this strain with the chimeras with pRS305, after linearizing the plasmid with *Bst*EII. We sporulated the correct transformants to obtain Nourseothricin resistant and Leucine prototroph segregants. We linearized the plasmids based on pRS306 by *Stu*I before transformation. To visualize centromere 4 segregation, first we built a diploid strain which is homozygous for *CENIV*-*tetO*, *tetR*-GFP, and *SPC29*-mCherry but heterozygous for *SPC105* (AJY5160). We digested pAJ817 (WT) and pAJ818 (BPM) by *Sac*II-*Kpn*I before transforming them in this strain. After that we sporulated the correct transformants to obtain the haploid strains.

We obtained the pSB148 from Biggins lab. In this plasmid, *IPL1* ORF is flanked by 1.0 Kb upstream as promoter (*prIPL1*) and 654 bp downstream sequence as terminator (*trIPL1*). We performed site directed mutagenesis with the Quick change kit (Agilant Technologies) to introduce the *ipl1-2* (H352Y) mutation in the *IPL1* ORF. We acquired *sgo1Δ* and *rts1Δ* from the yeast deletion library (Giaever et al., 2002). The strains with *sgo1Δ* demonstrated variable growth. Hence, we backcrossed them with YEF473 to remove any background mutations. We tested both normal and slow growing segregants for growth assays on YPD and YPD+benomyl. We confirmed the strains with bub1-ΔK (kinase deleted bub1) by western blot by anti-mCherry antibody and localization at unattached kinetochores.

To build the strains containing *ndc80-6A* or *dam1-3A*, first we deleted one allele of *NDC80* or *DAM1* ORF with *TRP1* marker. Next, we replaced the deleted allele with ndc80-6A (*KAN*) or dam1-3A (*KAN*) by transforming the transforming the heterozygous deletion strains of *NDC80* and *DAM1* with *Sac*II-*Apa*I digest of pAJ108 (ndc80-6A) and *Sac*II-*Sma*I digest of pSB617 (dam1-3A) respectively. After confirming the integration of the cassettes, we sporulated the diploids to obtain tryptophan auxotroph and G418 resistant segregants. We also confirmed the haploid strains with *ndc80-6A* and *dam1-3A* by colony check PCR followed by Sanger sequencing.

To construct the plasmids containing 6XHIS-MBP-Spc105^2-455, 222::GFP^, we used a pET28a chimera where MBP is already cloned within *Nhe*I-*Bam*HI sites. 6XHIS and MBP are separated by a linker of 13 amino acids (SSGLVPRGSHMAS) and a TEV protease cleavage site is present at the downstream of MBP. We cloned fragments containing SPC105^2-455, 222::GFP^ of WT and BPM within *Sal*I-*Eag*I sites, that are present at the downstream of MBP-TEV. Hence translation of the whole chimera will result into expression of a 1123 amino acids long protein in which 6XHIS-MBP-TEV is linked with the phosphodomain by a 9 amino acids long peptide fragment (GSEFELRRP).

### Phosphorylation of the N-terminus of Spc105

The post-translational modifications of Spc105 are cataloged in the pSYT database. The entry for Spc105 within this database can be viewed at the Global Protein Machine Database (Craig et al., 2004) using the following url: http://psyt.thegpm.org/~/dblist_pep_modmass/label=YGL093W&modmass=80@STY&display=0 Citations to the studies that observed the phosphorylated peptides are included in the main text.

### Cell culture

We grew yeast strains in YPD (yeast extract 1%, peptone 2%, dextrose 2%) at room temperature (25°C) and 32°C. For strains with W303 background, we supplemented the media with 0.1 mg/ml adenine. They were imaged in room temperature in presence of synthetic dextrose media, supplemented with essential amino acids. We added nocodazole and methionine to the mounting media to image the nocodazole arrested cells and Cdc20 depleted cells respectively. To form zygotes for constructing diploid strains, we mixed overnight grown cultures of two strains of opposite mating types and spotted on YPD plate which was incubated for 3-4 hours at 32°C. To induce sporulation, we grew diploid yeast cells were grown in YPD overnight to stationary phase. Then the cells were pelleted down and resuspended with starvation media (yeast extract 0.1%, Potassium acetate 1%) and incubated 4-5 days at RT.

Metaphase arrest by Cdc20 depletion: We synchronized the strains expressing Cdc20 from *prMET3* by treating them with α factor (2μg/ml) for 2 h in synthetic dextrose media lacking methionine. After that, they were washed to remove α factors and released in YPD supplemented with 2M methionine to knock down Cdc20 expression for 1-2 h. To release from Cdc20 repression mediated metaphase arrest, we washed the cells with synthetic media lacking methionine and incubated them in the same.

### Kinetochore particle pull down experiments

We used strains expressing Dsn1-His-Flag, Glc7-3XGFP, and either Spc105^222::mcherry^ or Spc105^BPM, 222::mCherry^. We purified the kinetochores as previously described (Gupta et al., 2017). Briefly, we harvested and lysed cells by blender in presence of liquid nitrogen and prepared clear lysate by ultracentrifugation at 24,000 r.p.m for 90min at 4°C. Equal amounts of precleared lysates were incubated with α-Flag-M2 conjugated magnetic dynabeads at 4°C for 2h 30min. The beads were then washed, and Flag-tagged protein was eluted from the beads by incubation with 0.5mg/ml 3xFlag peptide solution at room temperature for 30min. Western blotting assay was performed using commercial antibodies (α-Flag M2 (Sigma-Aldrich) 1:5000; α-Ds-Red (Santa Cruz Biotchnologies) 1:2000; α-GFP, JL-8 (Living Colors) 1:3000).

### Treatment of cells to study spindle pole and centromere IV (*CENIV*) separation

For this experiment, we used strains with TetO-CENIV, TetR-GFP, and Spc29-mCherry. For assay mentioned in Fig. 1C, we synchronized them at G1 stage with α factors (2 μg/ml) for 1h 45 min. They were then washed and released in YPD and cell samples were collected at 30, 45 and 60min. For assay mentioned in Fig. 4A, we utilized the strains of 5267 (Spc105^WT^, *sgo1Δ*) and 5273 (Spc105^BPM^, *sgo1Δ*) and synchronized them at G1 stage with α factors (2 μg/ml) for 1h 45 min. They were then washed and released in nocodazole supplemented (15 μg/ml) media to disrupt the spindle and activate the spindle checkpoint for 2 h. After that, we again washed them to remove nocodazole and release them in fresh YPD. Cell samples were collected and imaged at 0, 30 and 45 mins.

### Benomyl sensitivity assay

We prepared 10-fold serial dilutions of log-phase cultures starting from 0.1 OD_600_ and spotted on YPD or YPD containing 20 and /or 30 μg/ml benomyl (YPD+Ben^20^ and YPD+Ben^30^ respectively). Plates were incubated at 32°C and plate images were taken after 2 (on YPD) or 3 (YPD+benomyl) days. All benomyl spotting assays were performed with at least two or three biological replicates wherever possible and two technical replicates on both YPD+Ben^20^ and YPD+Ben^30^. This was done for the following reason. Benomyl is not readily soluble in media and it also has relatively poor thermal stability. Both these factors are problematic when making agar media, because benomyl must be dissolved in the agar once it reaches ∼ 60 °C. Because of this, we find that the effect of benomyl in YPD plates on control strains varies slightly from one batch to other. Therefore, we perform each spotting experiment at two different benomyl concentration. Furthermore, we included a wild-type positive control which should grow and a negative control strain which should either grow poorly or not grow at all in YPD+benomyl. If the strains behave as expected at both benomyl concentrations, we show the results from the higher concentration.

For the spotting assay shown in fig 4F, we incubated the ben^30^ plate for 5 days at 30°C as the growth rate of sgo1Δ and mad2Δ double mutant is poorer than *sgo1Δ* single mutant even in YPD. One should compare the growth of the double mutant of *mad2Δ* and bub1-ΔK (top panel) or the double mutant of *mad2Δ* and *sgo1Δ* (bottom panel) in the background of WT and BPM or RASA in benomyl containing media.

### Microscopy and image acquisition and analyses

A Nikon Ti-E inverted microscope with a 1.4 NA, 100X, oil-immersion objective was used in microscopy (Joglekar et al., 2013). A ten-plane Z-stack was acquired (200nm separation between adjacent planes). To measure Bub3 and Mad1-mCherry, the 1.5x opto-var lens was used. Total fluorescence of kinetochore clusters (16 kinetochores in metaphase) was measured by integrating the intensities over a 6×6 region centered on the maximum intensity pixel. The median intensity of pixels immediately surrounding the 6×6 area was used to correct for background fluorescence. The calculations were performed using ImageJ or semi-automated MATLAB programs as described earlier (Joglekar et al., 2006).

Analysis of kinetochore biorientation was conducted using a MATLAB GUI according to the scheme developed previously (Marco et al., 2013). Briefly, a region of interest containing the two spindle poles were manually selected from each cell. The script then identified the locations of the spindle poles and rotated the image so that the spindle axis (defined by the two spindle poles) coincides with the horizontal axis. Next, the fluorescence intensity in the GFP channel (used to record Spc105^222∷GFP^) over a rectangular region centered on the spindle axis and extending 4 pixels on either side of the spindle axis was summed to define the kinetochore distribution at each pixel along the spindle axis. The kinetochore distribution was then normalized by the total fluorescence to obtain the fluorescence line scan for each cell. Line-scans from > 50 cells were then displayed as a heat map in Fig. 1D and 4D. G1-arrested yeast cells expressing either wild-type Spc105 or Spc105^BPM^ were released into the cell cycle, and the distribution of kinetochores along the spindle axis (defined by the locations of the two spindle poles) was analyzed at 15-minute time intervals and visualized as a heat-map. Because yeast cells enter mitosis after ∼ 30 minutes following release from a G1 arrest, only 45 and 60-minute time points are shown in Fig. 1D and 45-minute time point is shown in 4D. In this visualization, each pixel row represents the kinetochore fluorescence line scan for a single cell, with the pixel rows have been sorted according to spindle length (longest to shortest from top to bottom, > 50 cells for each cell for each time-point).

Quantitative assessment of the relative distribution of kinetochores in the two spindle halves was obtained on the basis of the normalized kinetochore distribution as calculated above. Briefly, the brightest pixel in each spindle half was identified from the fluorescence line scan, and the absolute value of the intensity difference between these pixels was used as the measure of asymmetry in kinetochore distribution.

### Statistical analyses

To quantify the frequency with which metaphase cells with visibly recruited SAC proteins (Bub1, Bub3, or Mad1) at bioriented kinetochores in a cell population, we performed imaging each strain at least twice to obtain significant number of metaphase cells (>50). The scoring analysis divided the metaphase cell population into two groups: cells with visible SAC protein localization and the cells without detectable localization. The variation in the fraction of cells with visible SAC protein localization from individual experiments did not vary significantly. Therefore, we pooled observations from all experiments. To ascertain the statistical significance of these categorical scoring data, we applied Fisher’s exact test in Graphpad Prism (version 7). The number of cells analyzed for each strain is noted in the figure legends. To ascertain the statistical significance of the rest of the data, we applied the two-way ANOVA test using Graphpad Prism. The p-values from these tests are noted in the figure legends.

### Flow cytometry

We performed flow cytometry as described previously (Baum and Clarke, 2000). Starting from overnight inoculum, we grew the yeast strains to mid log phase. Then we added nocodazole (final concentration 15μg/ml) to the cultures to depolymerize spindle and activate the spindle checkpoint (Gillett et al., 2004). We collected the cell samples of 0.1 OD_600_ at designated time points (0, 1, 2 and 3 h).

### Spc105 phosphodomain Purification

The 6xHIS-MBP-Spc105^2-455, 222::GFP^ fusion construct was expressed in BL21-Rosetta 2 (DE3) (Novagen) or T7 expression *E. coli* cells (NEB) using 0.25 mM IPTG for 16 hours at 18 °C. Cells were then harvested, and resuspended in the lysis buffer (20 mM Tris (pH 7.5), 300 mM NaCl, 5 mM Imidazole, 0.1% TritonX-100 (Sigma), 5% glycerol, 1 mM PMSF) supplemented with complete EDTA-free protease inhibitor mix (Roche). Cells were lysed by sonication (3X for 3 minutes with 30 second pulse on and 30 second pulse off). Cleared supernatant was incubated with Ni-NTA agarose beads (Invitrogen) for 2 h at 4°C. Beads were washed with lysis buffer containing 500mM NaCl and 30mM Imidazole, and protein was eluted with lysis buffer containing 150mM NaCl and 250mM Imidazole. Subsequently, the protein was loaded onto a 16/60 Superdex 200 size exclusion column (GE) equilibrated with gel filtration buffer (20 mM Tris pH 7.5, 100 mM NaCl, 5% glycerol, and 1 mM DTT). Fractions were analyzed by SDS-PAGE and Coomassie staining, and peak fractions were aliquoted and stored at −80 °C.

### Preparation of X-rhodamine labeled microtubules

X-rhodamine (Cytoskeleton) labeled microtubule was achieved by using a mixture of X-rhodamine (1.25mg/ml) and unlabeled tubulins(10mg/ml) in BRB80 buffer (80 mM PIPES/KOH, pH 6.8, 1 mM MgCl_2_, 1 mM EGTA) containing 1 mM GTP and 4 mM MgCl_2_ at 37 °C. Microtubules were further stabilized by the addition of 10 μM Taxol and incubation at 37 °C for 15 min. Microtubules were pelleted down by ultracentrifugation (55,000 rpm for 10 minutes at 37°C) to get rid of unpolymerized tubulin.

### Decoration of Polystyrene beads with recombinant Spc105 phosphodomain

The assay was performed as previously described with small modifications (Espeut et al., 2012). In brief, decoration of Spc105 proteins on polystyrene beads was achieved using streptavidin-biotin system. 100 nm of WT and mutant Spc105 proteins was incubated with 10 µl of 0.1% w/v streptavidin polystyrene beads (Spherotech) conjugated with anti-penta-his biotin antibody (Qiagen) for 1 and half hours at 4°C. Before imaging, the beads were sonicated for 3 minutes in a water bath containing ice cubes. Subsequently, the beads were introduced inside the flow chamber coated with taxol stabilized microtubules. The images were acquired using TIRF microscope.

### Sample preparation for TIRF imaging

Flow cells were created as described before (Verma et al., 2015). Flow cells were first incubated with 30 μl of 1:100 dilution of monoclonal anti-β-Tubulin antibody (Sigma) for 10 minutes in a humidified chamber, followed by another incubation of 0.5% w/v Pluronic F-127 (Sigma) for 10 minutes, and 30 μl of X-rhodamine labeled microtubule for 20 minutes. The flow cells were then incubated with 30 μl of 2.5mg/ml casein for 5 minutes to block nonspecific sites. After blocking, flow cells were incubated with 30 μl of polystyrene beads decorated with WT or mutant Spc105 proteins and scavenging system (40 nM _D_-glucose, 250nM glucose oxidase, 64nM catalase and 1% (v/v) β-mercaptoethanol in BRB80 containing 50 mM NaCl (80 mM PIPES, 1 mM MgCl_2_, 1 mM EGTA, 50mM NaCl, pH 6.8). Subsequently, chambers were sealed, and samples were immediately imaged using TIRF microscope.

### TIRF microscopy

All images were acquired as described before (Verma V et. al, Plos one, 2015) with little modifications. In brief, Images were acquired on a Nikon Ti-E microscope equipped with a 100X 1.4 NA CFI-Apo oil immersion objective, an EMCCD camera (iXon+ DU 897; Andor), a 3 line (488, 561 and 640nm) monolithic laser combiner with AOTF laser system (Agilent) and Nikon NIS-Elements software. Images of X-rhodamine labeled tubulin and Spc105-GFP were acquired at the following settings: 600 frames @ 50ms exposure time, conversion gain-1X, EM multiplier gain setting 288, 561 laser power-40% and 488 laser power-40%. For TIRF image analysis, length of the microtubule was calculated with the help of Fiji software, and intensity of the Spc105-GFP conjugated with streptavidin polystyrene beads was measured using a custom built software in MATLAB as described before (Joglekar et al., 2006).

### Artificial tethering of Glc7 to the N-terminus of Spc105

For these set of experiments, we used the yeast strains where Glc7-Fkbp can be loaded on N-termini of Spc105 (FRB-GFP-Spc105) by adding rapamycin to the culture. To investigate Glc7 loading in prophase, haploid strains of were arrested in G1 by α factor treatment (2 μg/ml) for 1h 45 min. They were then washed and released in YPD media supplemented with methionine (to repress *CDC20* expression and block anaphase onset). We added nocodazole to the media (15 μg/ml) to disrupt the spindle structure and activate the spindle assembly checkpoint at 30 min after release from the G1 arrest (time at which cells are in S-phase with duplicated spindle pole bodies, and are in the process of kinetochore biorientation),. After 30 min., the cells were washed again to remove nocodazole, and released in rapamycin and methionine supplemented growth media for 30mins before imaging. To load Glc7 in metaphase, we repressed *CDC20* expression for 1h 10min. Then we supplemented the media with rapamycin and incubated for 30min before imaging.

**Figure S1.**
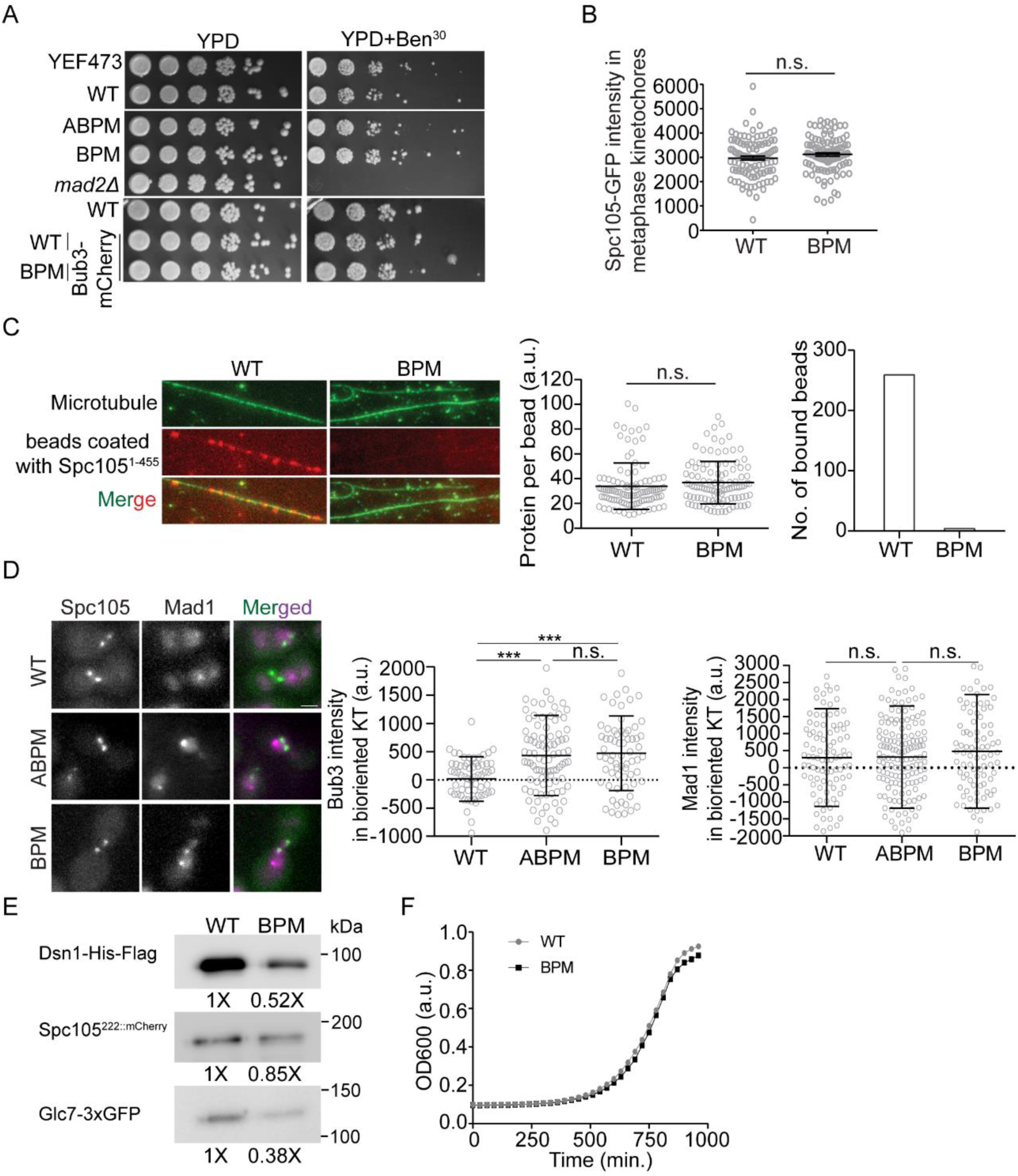
Related to Fig. 1 and 2. **(A)** Serial dilutions of yeast cells of the indicated genotype on rich media (YPD) and media containing Benomyl. YEF473 is the parent strain for nearly all the strains constructed in this study. WT refers to a wild-type strain expressing Spc105^222:GFP^ (GFP inserted at amino acid 222 in Spc105). **(B)** Quantification of the fluorescence due to Spc105^222:GFP^ and Spc105^222:GFP, BPM^ in metaphase kinetochore clusters (p=0.1605 using t-test). **(C)** Left panel: Total Internal Reflection Fluorescence micrographs showing the interaction of polystyrene beads coated with recombinant Spc105 phosphodomain (red, amino acid residues 1-455, containing either wild-type or mutant basic patches) with X-rhodamine labelled, Taxol-stabilized microtubules (green). Middle panel: Quantification of the average amount of protein conjugated per bead (n = 108 from 2 experiments, p=0.24 using the unpaired t-test). Right panel: Number of interactions observed for the respective phosphodomain (n = 259 and 4 for WT and BPM respectively from 2 experiments). **(D)** Representative images of metaphase cells expressing the indicated protein. Scatter plots: Quantification of Bub3-mCherry and Mad1-mCherry recruitment by bioriented kinetochores in metaphase-arrested cells of the indicated genotype. The data are presented as mean+ s.d. (n = 62., 95, 68, for WT, ABPM and BPM respectively pooled from 2 experiments in Bub3-mCherry intensity measurement, and n=103, 165 and 100 for WT, ABPM and BPM respectively pooled from 2 assays in Mad1-mCherry intensity measurement; p<0.0001 and 0.1622 for Bub3 and Mad1 respectively using t-test). Scale bar∼3.2μm. **(E)** Replicate data related to Figure 2A. Here, the loading volumes were adjusted to equalize the amount of Spc105 co-precipitating with Dsn1-Flag to make it possible to visually assess the amount Glc7-3xGFP co-precipitation. **(F)** Graph depicting the optical density measurements of strains expressing Spc105^WT^ and Spc105^BPM^ at the indicated time points as measured by 96-well plate reader.

**Figure S2.**
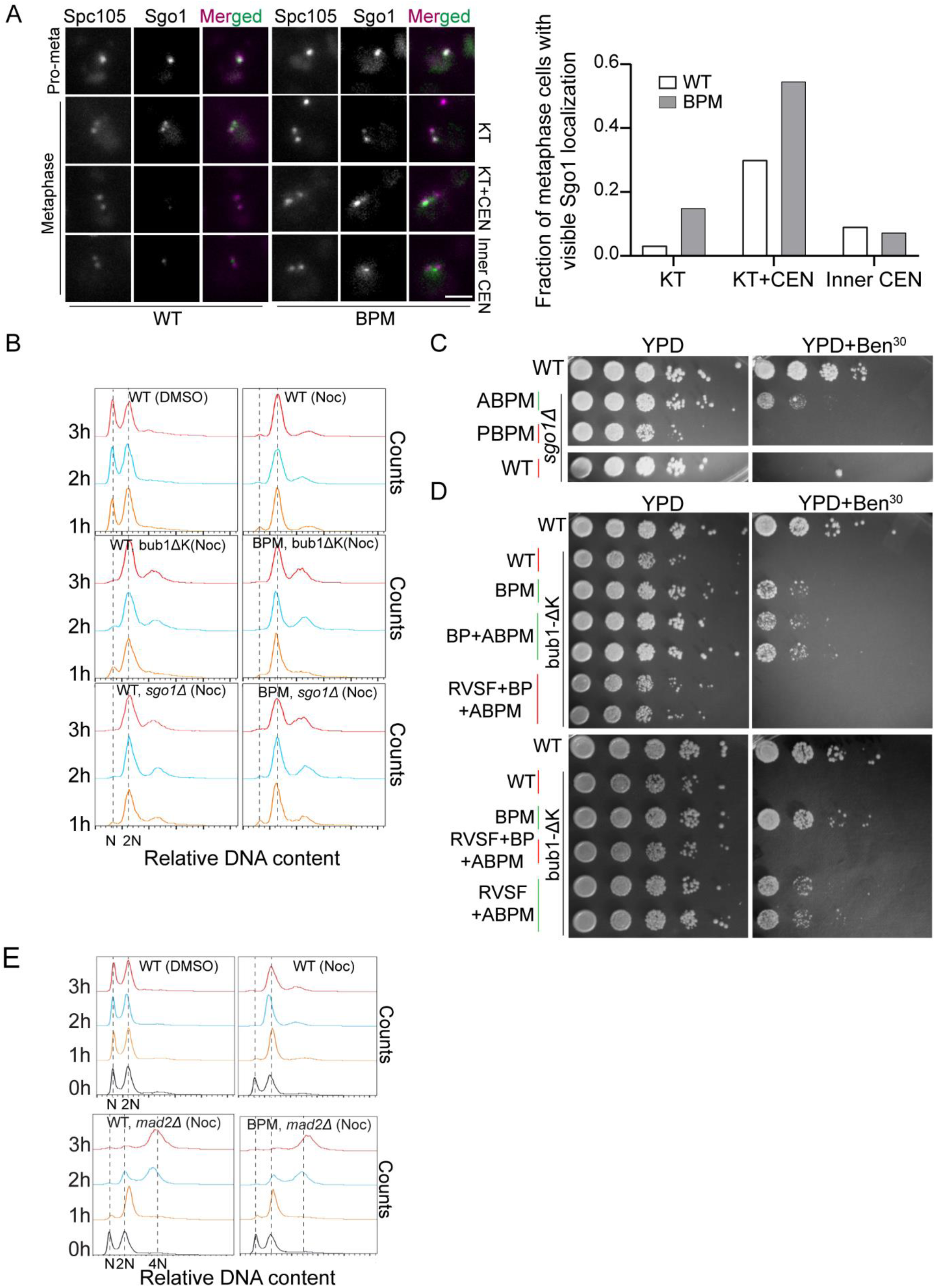
Related to Fig. 3 and 4. **(A)** Left panel: Representative micrographs showing the localization of Sgo1-GFP and Spc105^222:mCherry^ in prometaphase (single kinetochore cluster) and metaphase (two distinct kinetochore clusters) cells. The designations used for scoring cell populations is indicated on the right hand side (KT = kinetochore-colocalized, KT+CEN = Sgo1 colocalizes with both kinetochores and the centromeric region in between the two kinetochore clusters, and Inner CEN = Sgo1 localized mainly in the centromeric region, positioned between two sister kinetochore foci). Scale bar∼3.2μm. Right panel: Scoring shown in the bar graph on the right. The numbers mentioned above, represents the number of cells we examined for this assay. (n = 67 and 209 for WT and BPM respectively pooled from 2 experiments, p value according to Fisher exact test: <0.0084 for KT and KT+CEN and 0.6037 for inner CEN). **(B)** Flow cytometry-based evaluation of SAC signaling activity in strains lacking Sgo1 or that express Bub1^Δkinase^. **(C-D)** Benomyl spotting assay of the indicated strains. **(E)** Quantification of DNA content using flow cytometry after activation of the SAC by nocodazole treatment (0h). The first peak in each graph corresponds to cells with 1n DNA content (G1), whereas the second peak corresponds to cells with 2n DNA content (G2/M). In cells with an inactive SAC (*mad2Δ*), cells fail to arrest in M, and shift to the 4n peak (tetraploid).

**Figure S3.**
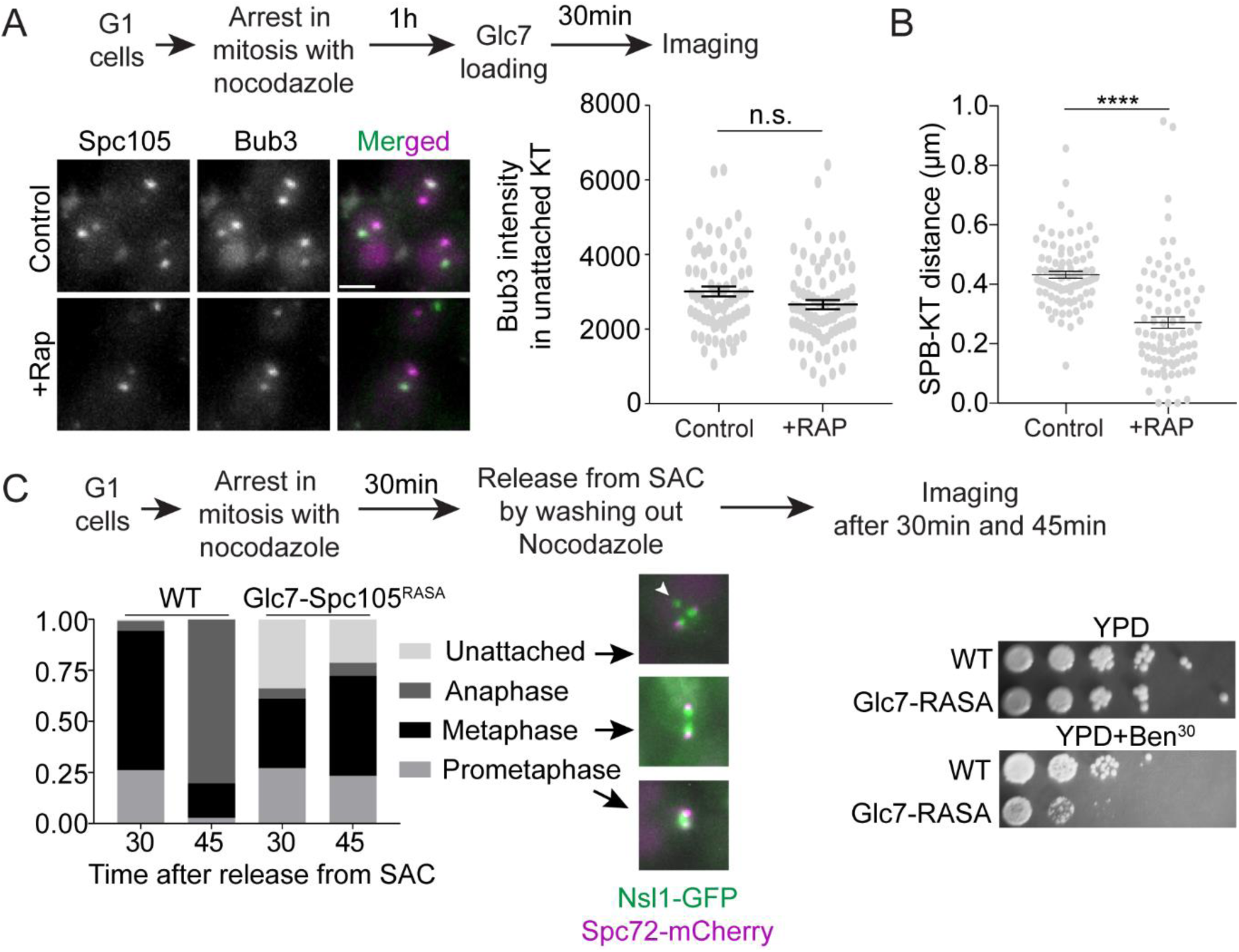
Related to Fig. 5. **(A)** Top: Flow chart describes the work-flow to tether Glc7 to the N-terminus of Spc105 in presence of nocodazole. Bottom left: Representative images of FRB-GFP-Spc105, Bub3-mCherry and merged in unattached kinetochores of control and rapamycin treated cells. Scale bar∼3.2μm. Bottom right: Scatter plot showing Bub3-mCherry intensities in unattached kinetochores of rapamycin treated and untreated control cells (mean± s.e.m., p=0.0585 obtained from unpaired t-test). The number of kinetochores analyzed for this assay: n = 71 and 78 for control and rapamycin treated cells respectively pooled from 2 experiments performed using two biological replicates. **(B)** Scatter plot showing SPB-KT distances in untreated control and rapamycin-treated cells. The data is presented as mean ± s.e.m. (n = 87 control and rapamycin treated cells respectively, data were pooled from three experimental repeats. p<0.0001 with unpaired t-test) **(C)** Comparative characterization of the kinetics of kinetochore biorientation in the strain expressing the chimeric molecule Glc7-Spc105^RASA^ (constructed by Rosenberg et al., 2011, work flow shown at the top). Bottom left: Bar graph displays the frequency of the indicated phenotypes. The number of number of kinetochores analyzed for this assay: n = 122 and 118 for WT and Glc7-RASA at 30min, 118 and 94 for WT and Glc7-RASA at 45min pooled from 2 experiments performed using two biological replicates. Bottom right: spotting assay indicates that the *GLC7-spc105^RASA^* strain is sensitive to benomyl.

## References

1. Akiyoshi, B., C.R. Nelson, J.A. Ranish, and S. Biggins. 2009. Analysis of Ipl1-Mediated Phosphorylation of the Ndc80 Kinetochore Protein in Saccharomyces cerevisiae. Genetics. 183:1591–1595.

2. Akiyoshi, B., K.K. Sarangapani, A.F. Powers, C.R. Nelson, S.L. Reichow, H. Arellano-Santoyo, T. Gonen, J.A. Ranish, C.L. Asbury, and S. Biggins. 2010. Tension directly stabilizes reconstituted kinetochore-microtubule attachments. Nature. 468:576–579.

3. Aravamudhan, P., R. Chen, B. Roy, J. Sim, and A.P. Joglekar. 2016. Dual mechanisms regulate the recruitment of spindle assembly checkpoint proteins to the budding yeast kinetochore. Mol Biol Cell. 27:3405–3417.

4. Aravamudhan, P., I. Felzer-Kim, K. Gurunathan, and A.P. Joglekar. 2014. Assembling the protein architecture of the budding yeast kinetochore-microtubule attachment using FRET. Curr Biol. 24:1437–1446.

5. Aravamudhan, P., A.A. Goldfarb, and A.P. Joglekar. 2015. The kinetochore encodes a mechanical switch to disrupt spindle assembly checkpoint signalling. Nat Cell Biol. 17:868–879.

6. Ayaz, P., X. Ye, P. Huddleston, C.A. Brautigam, and L.M. Rice. 2012. A TOG:alphabeta-tubulin complex structure reveals conformation-based mechanisms for a microtubule polymerase. Science. 337:857–860.

7. Bajaj, R., M. Bollen, W. Peti, and R. Page. 2018. KNL1 Binding to PP1 and Microtubules Is Mutually Exclusive. Structure. 26:1327–1336 e1324.

8. Baum, M., and L. Clarke. 2000. Fission Yeast Homologs of Human CENP-B Have Redundant Functions Affecting Cell Growth and Chromosome Segregation. Molecular and Cellular Biology. 20:2852–2864.

9. Bouck, D.C., and K. Bloom. 2007. Pericentric chromatin is an elastic component of the mitotic spindle. Curr Biol. 17:741–748.

10. Cheeseman, I.M., S. Anderson, M. Jwa, E.M. Green, J. Kang, J.R. Yates, 3rd, C.S. Chan, D.G. Drubin, and G. Barnes. 2002. Phospho-regulation of kinetochore-microtubule attachments by the Aurora kinase Ipl1p. Cell. 111:163–172.

11. Craig, R., J.P. Cortens, and R.C. Beavis. 2004. Open source system for analyzing, validating, and storing protein identification data. J Proteome Res. 3:1234–1242.

12. Espeut, J., D.K. Cheerambathur, L. Krenning, K. Oegema, and A. Desai. 2012. Microtubule binding by KNL-1 contributes to spindle checkpoint silencing at the kinetochore. J. Cell Biol. 196:469–482.

13. Etemad, B., T.E. Kuijt, and G.J. Kops. 2015. Kinetochore-microtubule attachment is sufficient to satisfy the human spindle assembly checkpoint. Nat Commun. 6:8987.

14. Franck, A.D., A.F. Powers, D.R. Gestaut, T. Gonen, T.N. Davis, and C.L. Asbury. 2007. Tension applied through the Dam1 complex promotes microtubule elongation providing a direct mechanism for length control in mitosis. Nat Cell Biol. 9:832–837.

15. Giaever, G., A.M. Chu, L. Ni, C. Connelly, L. Riles, S. Véronneau, S. Dow, A. Lucau-Danila, K. Anderson, B. André, A.P. Arkin, A. Astromoff, M. El Bakkoury, R. Bangham, R. Benito, S. Brachat, S. Campanaro, M. Curtiss, K. Davis, A. Deutschbauer, K.-D. Entian, P. Flaherty, F. Foury, D.J. Garfinkel, M. Gerstein, D. Gotte, U. Güldener, J.H. Hegemann, S. Hempel, Z. Herman, D.F. Jaramillo, D.E. Kelly, S.L. Kelly, P. Kötter, D. LaBonte, D.C. Lamb, N. Lan, H. Liang, H. Liao, L. Liu, C. Luo, M. Lussier, R. Mao, P. Menard, S.L. Ooi, J.L. Revuelta, C.J. Roberts, M. Rose, P. Ross-Macdonald, B. Scherens, G. Schimmack, B. Shafer, D.D. Shoemaker, S. Sookhai-Mahadeo, R.K. Storms, J.N. Strathern, G. Valle, M. Voet, G. Volckaert, C.-y. Wang, T.R. Ward, J. Wilhelmy, E.A. Winzeler, Y. Yang, G. Yen, E. Youngman, K. Yu, H. Bussey, J.D. Boeke, M. Snyder, P. Philippsen, R.W. Davis, and M. Johnston. 2002. Functional profiling of the Saccharomyces cerevisiae genome. Nature. 418:387.

16. Gillett, E.S., C.W. Espelin, and P.K. Sorger. 2004. Spindle checkpoint proteins and chromosome– microtubule attachment in budding yeast. The Journal of Cell Biology. 164:535–546.

17. Gupta, A., R.K. Evans, L.B. Koch, A.J. Littleton, and S. Biggins. 2018. Purification of kinetochores from the budding yeast Saccharomyces cerevisiae. Methods in cell biology. 144:349–370.

18. Gupta, A., B. Sripa, and T. Tripathi. 2017. Purification and characterization of two-domain glutaredoxin in the parasitic helminth Fasciola gigantica. Parasitol. Int. 66:432–435.

19. Haruki, H., J. Nishikawa, and U.K. Laemmli. 2008. The anchor-away technique: rapid, conditional establishment of yeast mutant phenotypes. Mol Cell. 31:925–932.

20. Hendrickx, A., M. Beullens, H. Ceulemans, T. Den Abt, A. Van Eynde, E. Nicolaescu, B. Lesage, and M. Bollen. 2009. Docking motif-guided mapping of the interactome of protein phosphatase-1. Chem. Biol. 16:365–371.

21. Hiruma, Y., C. Sacristan, S.T. Pachis, A. Adamopoulos, T. Kuijt, M. Ubbink, E. von Castelmur, A. Perrakis, and G.J. Kops. 2015. CELL DIVISION CYCLE. Competition between MPS1 and microtubules at kinetochores regulates spindle checkpoint signaling. Science. 348:1264–1267.

22. Ji, Z., H. Gao, and H. Yu. 2015. CELL DIVISION CYCLE. Kinetochore attachment sensed by competitive Mps1 and microtubule binding to Ndc80C. Science. 348:1260–1264.

23. Joglekar, A., R. Chen, and J. Lawrimore. 2013. A Sensitized Emission Based Calibration of FRET Efficiency for Probing the Architecture of Macromolecular Machines. Cellular and Molecular Bioengineering. 6:369–382.

24. Joglekar, A.P., D.C. Bouck, J.N. Molk, K.S. Bloom, and E.D. Salmon. 2006. Molecular architecture of a kinetochore–microtubule attachment site. Nature Cell Biology. 8:581.

25. Jürg Bähler, Jian-Qiu Wu, Mark S. Longtine, Nirav G. Shah, Amos Mckenzie III, Alexander B. Steever, Achim Wach, Peter Philippsen, and J.R. Pringle. 1998. Heterologous modules for efficient and versatile PCR-based gene targeting in Schizosaccharomyces pombe. YEAST. 14:943–951.

26. Kanshin, E., S. Giguere, C. Jing, M. Tyers, and P. Thibault. 2017. Machine Learning of Global Phosphoproteomic Profiles Enables Discrimination of Direct versus Indirect Kinase Substrates. Mol. Cell. Proteomics. 16:786–798.

27. Kawashima, S.A., Y. Yamagishi, T. Honda, K. Ishiguro, and Y. Watanabe. 2010a. Phosphorylation of H2A by Bub1 prevents chromosomal instability through localizing shugoshin. Science. 327:172–177.

28. Kawashima, S.A., Y. Yamagishi, T. Honda, K.-i. Ishiguro, and Y. Watanabe. 2010b. Phosphorylation of H2A by Bub1 Prevents Chromosomal Instability Through Localizing Shugoshin. Science. 327:172–177.

29. Lampson, M.A., K. Renduchitala, A. Khodjakov, and T.M. Kapoor. 2004. Correcting improper chromosome-spindle attachments during cell division. Nat Cell Biol. 6:232–237.

30. Liu, D., M. Vleugel, C.B. Backer, T. Hori, T. Fukagawa, I.M. Cheeseman, and M.A. Lampson. 2010. Regulated targeting of protein phosphatase 1 to the outer kinetochore by KNL1 opposes Aurora B kinase. J. Cell Biol. 188:809–820.

31. London, N., S. Ceto, J.A. Ranish, and S. Biggins. 2012. Phosphoregulation of Spc105 by Mps1 and PP1 regulates Bub1 localization to kinetochores. Curr Biol. 22:900–906.

32. Marco, E., J.F. Dorn, P.H. Hsu, K. Jaqaman, P.K. Sorger, and G. Danuser. 2013. S. cerevisiae chromosomes biorient via gradual resolution of syntely between S phase and anaphase. Cell. 154:1127–1139.

33. Meadows, John C., Lindsey A. Shepperd, V. Vanoosthuyse, Theresa C. Lancaster, Alicja M. Sochaj, Graham J. Buttrick, Kevin G. Hardwick, and Jonathan B.A. Millar. 2011. Spindle Checkpoint Silencing Requires Association of PP1 to Both Spc7 and Kinesin-8 Motors. Developmental Cell. 20:739–750.

34. Munoz-Barrera, M., I. Aguilar, and F. Monje-Casas. 2015. Dispensability of the SAC Depends on the Time Window Required by Aurora B to Ensure Chromosome Biorientation. PLoS One. 10:e0144972.

35. Nijenhuis, W., G. Vallardi, A. Teixeira, G.J. Kops, and A.T. Saurin. 2014. Negative feedback at kinetochores underlies a responsive spindle checkpoint signal. Nat Cell Biol. 16:1257–1264.

36. Pearson, C.G., P.S. Maddox, T.R. Zarzar, E.D. Salmon, and K. Bloom. 2003. Yeast Kinetochores Do Not Stabilize Stu2p-dependent Spindle Microtubule Dynamics. Molecular Biology of the Cell. 14:4181–4195.

37. Peplowska, K., A.U. Wallek, and Z. Storchova. 2014. Sgo1 regulates both condensin and Ipl1/Aurora B to promote chromosome biorientation. PLoS Genet. 10:e1004411.

38. Pinsky, B.A., C. Kung, K.M. Shokat, and S. Biggins. 2006. The Ipl1-Aurora protein kinase activates the spindle checkpoint by creating unattached kinetochores. Nat Cell Biol. 8:78–83.

39. Posch, M., G.A. Khoudoli, S. Swift, E.M. King, J.G. Deluca, and J.R. Swedlow. 2010. Sds22 regulates aurora B activity and microtubule-kinetochore interactions at mitosis. J. Cell Biol. 191:61–74.

40. Primorac, I., J.R. Weir, E. Chiroli, F. Gross, I. Hoffmann, S. van Gerwen, A. Ciliberto, and A. Musacchio. 2013. Bub3 reads phosphorylated MELT repeats to promote spindle assembly checkpoint signaling. Elife. 2:e01030.

41. Rosenberg, J.S., F.R. Cross, and H. Funabiki. 2011. KNL1/Spc105 recruits PP1 to silence the spindle assembly checkpoint. Curr Biol. 21:942–947.

42. Salic, A., J.C. Waters, and T.J. Mitchison. 2004. Vertebrate shugoshin links sister centromere cohesion and kinetochore microtubule stability in mitosis. Cell. 118:567–578.

43. Scott, R.J., C.P. Lusk, D.J. Dilworth, J.D. Aitchison, and R.W. Wozniak. 2005. Interactions between Mad1p and the nuclear transport machinery in the yeast Saccharomyces cerevisiae. Mol Biol Cell. 16:4362–4374.

44. Smolka, M.B., C.P. Albuquerque, S.-h. Chen, and H. Zhou. 2007. Proteome-wide identification of in vivo targets of DNA damage checkpoint kinases. Proceedings of the National Academy of Sciences. 104:10364–10369.

45. Suijkerbuijk, Saskia J.E., M. Vleugel, A. Teixeira, and Geert J.P.L. Kops. 2012. Integration of Kinase and Phosphatase Activities by BUBR1 Ensures Formation of Stable Kinetochore-Microtubule Attachments. Developmental Cell. 23:745–755.

46. Suzuki, A., A. Gupta, S.K. Long, R. Evans, B.L. Badger, E.D. Salmon, S. Biggins, and K. Bloom. 2018. A Kinesin-5, Cin8, Recruits Protein Phosphatase 1 to Kinetochores and Regulates Chromosome Segregation. Curr Biol. 28:2697–2704 e2693.

47. Tanaka, T.U., N. Rachidi, C. Janke, G. Pereira, M. Galova, E. Schiebel, M.J. Stark, and K. Nasmyth. 2002. Evidence that the Ipl1-Sli15 (Aurora kinase-INCENP) complex promotes chromosome bi-orientation by altering kinetochore-spindle pole connections. Cell. 108:317–329.

48. Tauchman, E.C., F.J. Boehm, and J.G. DeLuca. 2015. Stable kinetochore-microtubule attachment is sufficient to silence the spindle assembly checkpoint in human cells. Nat Commun. 6:10036.

49. Verma, V., L. Mallik, R.F. Hariadi, S. Sivaramakrishnan, G. Skiniotis, and A.P. Joglekar. 2015. Using Protein Dimers to Maximize the Protein Hybridization Efficiency with Multisite DNA Origami Scaffolds. PLoS One. 10:e0137125.

50. Verzijlbergen, K.F., O.O. Nerusheva, D. Kelly, A. Kerr, D. Clift, F. de Lima Alves, J. Rappsilber, and A.L. Marston. 2014. Shugoshin biases chromosomes for biorientation through condensin recruitment to the pericentromere. Elife. 3:e01374.

51. Welburn, J.P.I., M. Vleugel, D. Liu, J.R. Yates Iii, M.A. Lampson, T. Fukagawa, and I.M. Cheeseman. 2010. Aurora B Phosphorylates Spatially Distinct Targets to Differentially Regulate the Kinetochore-Microtubule Interface. Molecular Cell. 38:383–392.

52. Xu, Z., B. Cetin, M. Anger, U.S. Cho, W. Helmhart, K. Nasmyth, and W. Xu. 2009. Structure and function of the PP2A-shugoshin interaction. Mol Cell. 35:426–441.

53. Yang, Y., D. Tsuchiya, and S. Lacefield. 2015. Bub3 promotes Cdc20-dependent activation of the APC/C in S. cerevisiae. J. Cell Biol. 209:519–527.

54. Zhang, G., T. Lischetti, and J. Nilsson. 2014. A minimal number of MELT repeats supports all the functions of KNL1 in chromosome segregation. J Cell Sci. 127:871–884.

