## Supplementary material for "Minimization of a harmful cross-talk between mitotic checkpoint silencing and error correction": Table S1

Table S1: Strains used in this study, which were constructed in Joglekar lab:

| Strain (AJY#) | Genotype | Used in figure |
| --- | --- | --- |
| 5199 | *MAT*a*, trp1Δ63 leu2Δ-1 ura3-52 his3Δ200 lys2-8Δ1, TetR-GFP (LEU2); SPC29-mCherry-HIS3, TetO-CENIV (URA3), spc105Δ::TRP1::*Spc105 ^222::mCherry^ *(KAN)* | 1C |
| 5201 | *MAT*a*, trp1Δ63 leu2Δ-1 ura3-52 his3Δ200 lys2-8Δ1, TetR-GFP (LEU2); SPC29-mCherry-HIS3, TetO-CENIV (URA3), spc105Δ::TRP1::*Spc105 ^222::mCherry, 101-104, 340-340::AAAA^ *(KAN)* | 1C |
| 5078 | *MAT*a *trp1Δ63 ura3-52 his3Δ200 lys2-8Δ1, spc105Δ::NAT*, *leu2Δ-1::* Spc105 ^222::GFP^ (*LEU2*), *SPC97*-mcherry-*HYG*, *prMET3-CDC20 (URA3)* | 1D |
| 5080 | *MAT*a *trp1Δ63 ura3-52 his3Δ200 lys2-8Δ1, spc105Δ::NAT*, *leu2Δ-1::* Spc105 ^222::GFP, 101-104, 340-343::AAAA^ (*LEU2*), *SPC97*-mcherry- *HYG*, *prMET3-CDC20 (URA3)* | 1D |
| 3635 | *MAT*a *trp1Δ63 ura3-52 his3Δ200 lys2-8Δ1, spc105Δ::NAT, leu2Δ-1::* Spc105^222::GFP^ *(LEU2), BUB3*-mcherry-*HYG, prMET3-CDC20-URA3* | 1E, S1B, S1D |
| 3637 | *MAT*a *trp1Δ63 ura3-52 his3Δ200 lys2-8Δ1, spc105Δ::NAT, leu2Δ-1::* spc105^222::GFP, 101-104::AAAA^ *(LEU2), BUB3*-mcherry-*HYG, prMET3-CDC20 (URA3)* | 1E, S1D |
| 3638 | *MAT*a *trp1Δ63 ura3-52 his3Δ200 lys2-8Δ1, spc105Δ::NAT, leu2Δ-1::* Spc105^222::GFP, 101-104, 340-343::AAAA^ *(LEU2), BUB3*-mcherry-*HYG, prMET3-CDC20 (URA3)* | 1E, S1B, S1D |
| 3797 | *MAT*a *trp1Δ63 ura3-52 his3Δ200 lys2-8Δ1, spc105Δ::NAT, leu2Δ-1::* Spc105^222::GFP^ *(LEU2), MAD1*-mCherry-*HYG,* *nup60∆::TRP1, prMET3-CDC20 (URA3)* | 1E, S1D |
| 3798 | *MAT*a *trp1Δ63 ura3-52 his3Δ200 lys2-8Δ1, spc105Δ::NAT, leu2Δ-1::* Spc105^222::GFP, 101-104, 340-343::AAAA^ *(LEU2), MAD1*-mCherry-*HYG, nup60∆::TRP1, prMET3-CDC20 (URA3)* | 1E, S1D |
| 3939 | *MatA, trp1Δ63 ura3-52 his3Δ200 lys2-8Δ1, spc105Δ::NAT, Spc105 222::GFP (LEU2) 101-104::AAAA , nup60Δ::Trp1, Mad1-mcherry:hyg, prMET3-Cdc20 (URA3)* | 1E, S1D |
| 5176 | *MAT*a *trp1Δ63 ura3-52 his3Δ200 lys2-8Δ1, spc105Δ::NAT, leu2Δ-1::* BP(Spc105^79-145^)*-* Spc105^222::GFP^*^,^* ^101-104::AAAA^ *(LEU2), BUB3-mCherry-HYG* | 1Fi |
| 5177 | *MAT*α *trp1Δ63 ura3-52 his3Δ200 lys2-8Δ1, spc105Δ::NAT, leu2Δ-1::* BP(Spc105^79-145^)*-* Spc105^222::GFP^*^,^* ^101-104::AAAA^ *(LEU2), BUB3-mCherry-HYG* | 1Fi |
| 5384 | *MAT*α *trp1Δ63 ura3-52 his3Δ200 lys2-8Δ1, spc105Δ::NAT, leu2Δ-1::* EBP2(Spc105^55-145^)*-* Spc105^222::GFP^*^,^* ^101-104::AAAA^ *(LEU2), BUB3-mCherry-HYG* | 1Fi |
| 3606 | *MAT*a *trp1Δ63 ura3-52 his3Δ200 lys2-8Δ1, spc105Δ::NAT, leu2Δ-1::* Spc105 ^222::GFP^ *(LEU2), BUB3-mCherry-HYG* | 1Fi, 2B, S1A |
| 3627 | *MAT*a *trp1Δ63 ura3-52 his3Δ200 lys2-8Δ1, spc105Δ::NAT, leu2Δ-1::* Spc105 ^222::GFP, 101-104, 340-340::AAAA^ *(LEU2), BUB3-mCherry-HYG* | 1Fi, 2B, S1A |
| 4650 | *MAT*a *trp1Δ63 ura3-52 his3Δ200 lys2-8Δ1, spc105Δ::NAT, Spc105 222::GFP (LEU2), Bub1-ymCherry-Hyg* | 1Fii, 4B, 5C |
| 4651 | *MAT*α *trp1Δ63 ura3-52 his3Δ200 lys2-8Δ1, spc105Δ::NAT, Spc105 222::GFP (LEU2), Bub1-ymCherry-Hyg* | 1Fii, 4B, 5C |
| 4652 | *MAT*α *trp1Δ63 ura3-52 his3Δ200 lys2-8Δ1, spc105Δ::NAT, Spc105 ^101-104, 340-343::AAAA^ 222::GFP (LEU2), Bub1-ymCherry-Hyg* | 1Fii, 4B, 5C |
| 4653 | *MAT*a *trp1Δ63 ura3-52 his3Δ200 lys2-8Δ1, spc105Δ::NAT, Spc105 ^101-104, 340-343::AAAA^ 222::GFP (LEU2), Bub1-ymCherry-Hyg* | 1Fii, 4B, 5C |
| 5732 | *MATa, trp1Δ63 ura3-52 his3Δ200 lys2-8Δ1, spc105Δ::NAT, RVSF+BP (Spc105^55-145^)-Spc105^222::GFP 101-104^ (LEU2), BUB1-ymCherry-HYG* | 1Fii |
| 5733 | *MAT*α, *trp1Δ63 ura3-52 his3Δ200 lys2-8Δ1, spc105Δ::NAT, RVSF+BP (Spc105^55-145^)-Spc105^222::GFP 101-104^ (LEU2), BUB1-ymCherry-HYG* | 1Fii |
| 4214 | *MatA,* *trp1Δ63 ura3-52 his3Δ200 lys2-8Δ1, spc105Δ::NAT, TOG2-Spc105^222::GFP^ (LEU2), Bub3-mCherry-Hyg, prMET3-Cdc20 (URA3)* | 1G |
| 4215 | *Matα, trp1Δ63 ura3-52 his3Δ200 lys2-8Δ1, spc105Δ::NAT, TOG2-Spc105 222::GFP 101-104::AAAA (LEU2), Bub3-mCherry-Hyg, prMET3-Cdc20 (URA3)* | 1G |
| 5523 | *MATa, trp1Δ63 ura3-52 his3Δ200 leu2∆-1, can1, spc105Δ::NAT, SPC10 ^222::mCherry^ (LEU2), Glc7-3XGFP-HIS3, Dsn1-HIS-FLAG (URA3)* | 2A, S1E |
| 5524 | *MATa, trp1Δ63 ura3-52 his3Δ200 leu2∆-1, can1, spc105Δ::NAT, SPC105 ^101-104, 340-343::AAAA 222::mCherry^ (LEU2), Glc7-3XGFP-HIS3, Dsn1-HIS-FLAG (URA3)* | 2A, S1E |
| 3626 | *MAT*a *trp1Δ63 ura3-52 his3Δ200 lys2-8Δ1, spc105Δ::NAT, leu2Δ-1::* Spc105 ^222::GFP^*^,^* ^101-104::AAAA^ *(LEU2), BUB3-mCherry-HYG* | 2B |
| 3133 | *MAT*a *trp1Δ63 ura3-52 his3Δ200 lys2-8Δ1, spc105Δ::NAT, leu2Δ-1:: Spc105* ^222::GFP^ *(LEU2)* | S1A, S1F, 2C-D, 3i-vi, 4C, 4E, 4F, 5D, S2B-E |
| 3448 | *MAT*a *trp1Δ63 ura3-52 his3Δ200 lys2-8Δ1, spc105Δ::NAT, leu2Δ-1::* Spc105 ^222::GFP^*^,^* ^101-104, 340-340::AAAA^ *(LEU2)* | S1A, S1F, 2C-D, 3i-iii, v-vi |
| 4384 | *MAT*a*, trp1Δ63 ura3-52 his3Δ200 lys2-8Δ1, spc105Δ::NAT, leu2Δ-1::* Spc105 ^222::GFP, 101-104, 340-340::AAAA^ *(LEU2),* bub1ΔK*-mCherry-HYG* | 3i, S2D, 4C, S2B |
| 4385 | *MAT*a*, trp1Δ63 ura3-52 his3Δ200 lys2-8Δ1, spc105Δ::NAT, leu2Δ-1::* Spc105 ^222::GFP, 101-104, 340-340::AAAA^ *(LEU2),* bub1ΔK*-mCherry-HYG* | 3i |
| 4415 | *MAT*a*, trp1Δ63 ura3-52 his3Δ200 lys2-8Δ1, spc105Δ::NAT, leu2Δ-1::* Spc105 ^222::GFP^ *(LEU2),* bub1ΔK*-mCherry-HYG* | 3i, 4C, 4F, S2D, S2B |
| 4416 | *MAT*a*, trp1Δ63 ura3-52 his3Δ200 lys2-8Δ1, spc105Δ::NAT, leu2Δ-1::* Spc105 ^222::GFP^ *(LEU2),* bub1ΔK*-mCherry-HYG* | 3i |
| 4485 | *MAT*a*, trp1Δ63 leu2Δ-1 ura3-52 his3Δ200 lys2-8Δ1, BUB1-*mCherry-*HYG* | 3i |
| 4203 | *MATα, spc105Δ::NAT, leu2Δ-1::* Spc105 ^222::^*^GFP^ (LEU2), sgo1Δ::Kan* | 3ii, 4F, 5D, S2B |
| 4204 | *MATα, spc105Δ::NAT, leu2Δ-1::* Spc105 ^222::GFP^ *(LEU2), sgo1Δ::Kan* | 3ii |
| 4205 | *MAT*a *, spc105Δ::NAT, leu2Δ-1::* Spc105 ^222::GFP, 101-104, 340-340::AAAA^ *(LEU2), sgo1Δ::Kan* | 3ii. S2B |
| 4206 | *MAT*a*, spc105Δ::NAT, leu2Δ-1::* Spc105 ^222::GFP, 101-104, 340-340::AAAA^ *(LEU2), sgo1Δ::Kan* | 3ii |
| 4233 | *MAT*a*, spc105Δ::NAT, leu2Δ-1::* Spc105 ^222::GFP^ *(LEU2), rts1Δ::Kan* | 3iii |
| 4234 | *MAT*a*, spc105Δ::NAT, leu2Δ-1::* Spc105 ^222::GFP^ *(LEU2), rts1Δ::Kan* | 3iii |
| 4235 | *MAT*a*, trp1Δ63 leu2Δ-1 ura3-52 his3Δ200 lys2-8Δ1, spc105Δ::NAT, leu2Δ-1::* Spc105 ^222::GFP, 101-104, 340-340::AAAA^ *(LEU2), rts1Δ::Kan* | 3iii |
| 4236 | *MAT*a*, spc105Δ::NAT, leu2Δ-1::* Spc105 ^222::GFP, 101-104, 340-340::AAAA^ *(LEU2), trp1Δ63 leu2Δ-1 ura3-52 his3Δ200 lys2-8Δ1, rts1Δ::Kan* | 3iii |
| 4022 | *MAT*a*, trp1Δ63 ura3-52 his3Δ200 lys2-8Δ1, spc105Δ::NAT, leu2Δ-1::* Spc105*-*6A *(LEU2)* | 3ii-iii, iv |
| 4338 | *trp1Δ63 ura3-52 his3Δ200 lys2-8Δ1, spc105Δ::NAT, leu2Δ-1::* Spc105 ^222::GFP^ *(LEU2), ipl1Δ::TRP1, ipl1-2 (H352Y, CEN, URA3)* | 3iv |
| 4339 | *trp1Δ63 ura3-52 his3Δ200 lys2-8Δ1, spc105Δ::NAT, leu2Δ-1::* Spc105 ^222::GFP^ *(LEU2), ipl1Δ::TRP1, ipl1-2 (H352Y, CEN, URA3)* | 3iv |
| 4340 | *trp1Δ63 ura3-52 his3Δ200 lys2-8Δ1, spc105Δ::NAT, leu2Δ-1::* Spc105 ^222::GFP, 101-104, 340-340::AAAA^ *(LEU2), ipl1Δ::TRP1, ipl1-2 (H352Y, CEN, URA3)* | 3iv |
| 4341 | *trp1Δ63 ura3-52 his3Δ200 lys2-8Δ1, spc105Δ::NAT, leu2Δ-1::* Spc105 ^222::GFP, 101-104, 340-340::AAAA^ *(LEU2), ipl1Δ::TRP1, ipl1-2 (H352Y, CEN, URA3)* | 3iv |
| 4357 | *trp1Δ63 ura3-52 his3Δ200 lys2-8Δ1, spc105Δ::NAT, leu2Δ-1::* Spc105 ^222::GFP^ *(LEU2), ipl1Δ::TRP1, IPL1 (CEN, URA3)* | 3iv |
| 4362 | *trp1Δ63 ura3-52 his3Δ200 lys2-8Δ1, spc105Δ::NAT, leu2Δ-1::* Spc105 ^222::GFP, 101-104, 340-340::AAAA^ *(LEU2), ipl1Δ::TRP1, IPL1 (CEN, URA3)* | 3iv |
| 4712 | *MAT*a *trp1Δ63 ura3-52 his3Δ200 lys2-8Δ1, spc105Δ::NAT, leu2Δ-1::* Spc105 ^222::GFP^ *(LEU2), ndc80Δ:: TRP1::* ndc80-6A-*12MYC (KAN)* | 3v |
| 4713 | *Matα trp1Δ63 ura3-52 his3Δ200 lys2-8Δ1, spc105Δ::NAT, leu2Δ-1::* Spc105 ^222::GFP^ *(LEU2), ndc80Δ:: TRP1::* ndc80-6A-*12MYC (KAN)* | 3v |
| 4714 | *MAT*a *trp1Δ63 ura3-52 his3Δ200 lys2-8Δ1, spc105Δ::NAT, leu2Δ-1::* Spc105 ^222::GFP, 101-104, 340-340::AAAA^ *(LEU2), ndc80Δ:: TRP1::* ndc80-6A-*12MYC (KAN)* | 3v |
| 4715 | *MATα trp1Δ63 ura3-52 his3Δ200 lys2-8Δ1, spc105Δ::NAT, leu2Δ-1::* Spc105 ^222::GFP, 101-104, 340-340::AAAA^ *(LEU2), ndc80Δ:: TRP1::* ndc80-6A-*12MYC (KAN)* | 3v |
| 4920 | *MATα trp1Δ63 leu2Δ-1 ura3-52 his3Δ200 lys2-8Δ1, dam1Δ::TRP1::*dam1*(*S257A S265A S292A*) (KAN)* | 3vi, 5D |
| 4921 | *MATα trp1Δ63 leu2Δ-1 ura3-52 his3Δ200 lys2-8Δ1, dam1Δ::TRP1::*dam1*(*S257A S265A S292A*) (KAN)* | 3vi |
| 4904 | *MAT*a *trp1Δ63 ura3-52 his3Δ200 lys2-8Δ1, spc105Δ::NAT, leu2Δ-1::* Spc105 ^222::GFP, 101-104, 340-340::AAAA^ *(LEU2), dam1Δ::TRP1::*dam1*(*S257A S265A S292A*) (KAN)* | 3vi |
| 4905 | *MAT*a *trp1Δ63 ura3-52 his3Δ200 lys2-8Δ1, spc105Δ::NAT, leu2Δ-1::* Spc105 ^222::GFP, 101-104, 340-340::AAAA^ *(LEU2), dam1Δ::TRP1::*dam1*(*S257A S265A S292A*) (KAN)* | 3vi |
| 4964 | *MAT*a *trp1Δ63 ura3-52 his3Δ200 lys2-8Δ1, dam1Δ::TRP1::*dam1*(*S257A S265A S292A*) (KAN), ndc80Δ:: TRP1::* ndc80-6A-*12MYC (KAN), NSL1-GFP-HIS3* | 3vi |
| 4965 | *MAT*a *trp1Δ63 ura3-52 his3Δ200 lys2-8Δ1, dam1Δ::TRP1::*dam1*(*S257A S265A S292A*) (KAN), ndc80Δ:: TRP1::* ndc80-6A-*12MYC (KAN), NSL1-GFP-HIS3* | 3vi |
| 5000 | *MAT*a *trp1Δ63 ura3-52 his3Δ200 lys2-8Δ1, dam1Δ::TRP1::*dam1 *(*S257A S265A S292A*) (KAN), ndc80Δ:: TRP1::* ndc80-6A-*12MYC (KAN), NSL1-GFP-HIS3, spc105Δ::NAT, leu2Δ-1::* Spc105^222::mCherry, 101-104, 340-340::AAAA^ *(LEU2)* | 3vi |
| 5273 | *MAT*a*, trp1Δ63 leu2Δ-1 ura3-52 his3Δ200 lys2-8Δ1, TetR-GFP (LEU2); SPC29-mCherry-HIS3, TetO-CENIV (URA3), spc105Δ::TRP1::*Spc105 ^222::mCherry^ *(KAN), sgo1Δ::TRP1* | 4A |
| 5267 | *MAT*a*, trp1Δ63 leu2Δ-1 ura3-52 his3Δ200 lys2-8Δ1, TetR-GFP (LEU2); SPC29-mCherry-HIS3, TetO-CENIV (URA3), spc105Δ::TRP1::*Spc105 ^222::mCherry, 101-104, 340-340::AAAA^ *(KAN), sgo1Δ::TRP1* | 4A |
| 5756 | *MATa, trp1Δ63 leu2Δ-1 ura3-52 his3Δ200 lys2-8Δ1, spc105Δ::NAT, Spc105 ^222::GFP, 21-24 (GILK)::AAAA^ (LEU2), BUB1-mCherry-HYG* | 4B |
| 5757 | *MATa, trp1Δ63 leu2Δ-1 ura3-52 his3Δ200 lys2-8Δ1, spc105Δ::NAT, Spc105 ^222::GFP, 21-24 (GILK)::AAAA^ (LEU2), BUB1-mCherry-HYG* | 4B |
| 5778 | *MATα, trp1Δ63 leu2Δ-1 ura3-52 his3Δ200 lys2-8Δ1, spc105Δ::NAT, leu2:Spc105^222::GFP GILK:AAAA, 101-104::AAAA^ (LEU2), BUB1-mCherry-HYG* | 4B |
| 5825 | *MATa, trp1Δ63 leu2Δ-1 ura3-52 his3Δ200 lys2-8Δ1, spc105Δ::NAT, leu2:Spc105 ^222::GFP 21-24 (GILK)::AAAA, 101-104::AAAA^ (LEU2), BUB1-mCherry-HYG* | 4B |
| 5764 | *MATα, , trp1Δ63 leu2Δ-1 ura3-52 his3Δ200 lys2-8Δ1, spc105Δ::NAT, leu2:Spc105 ^222::GFP, 21-24 (GILK)::AAAA, 101-104::AAAA^ (LEU2), bub1-∆K-mCherry-HYG* | 4C |
| 5765 | *MATα, , trp1Δ63 leu2Δ-1 ura3-52 his3Δ200 lys2-8Δ1, spc105Δ::NAT, Spc105^222::GFP, 21-24 (GILK)::AAAA^ (LEU2), bub1ΔK-mCherry-HYG* | 4C |
| 5840 | *MATa, trp1Δ63 ura3-52 his3Δ200 lys2-8Δ1, spc105Δ::NAT, leu2Δ-1::Spc105^222::GFP^ (LEU2), Spc97-mCherry-HYG, mad3Δ::KAN* | 4D |
| 5841 | *MATa, trp1Δ63 ura3-52 his3Δ200 lys2-8Δ1, spc105Δ::NAT, leu2Δ-1::Spc105^222::GFP^ (LEU2), Spc97-mCherry-HYG, mad3Δ::KAN* | 4D |
| 5842 | *MATa, trp1Δ63 ura3-52 his3Δ200 lys2-8Δ1, spc105Δ::NAT, leu2Δ-1::Spc105^222::GFP,^* *^RASA (V76, F78::A)^ (LEU2), Spc72-mCherry-HYG, mad3Δ::URA3* | 4D |
| 5843 | *MATa, trp1Δ63 ura3-52 his3Δ200 lys2-8Δ1, spc105Δ::NAT, leu2Δ-1::Spc105^222::GFP,^* *^RASA (V76, F78::A)^ (LEU2), Spc72-mCherry-HYG, mad3Δ::URA3* | 4D |
| 4555 | *MAT*a *trp1Δ63 ura3-52 his3Δ200 lys2-8Δ1, spc105Δ::NAT, leu2Δ-1::* Spc105 ^222::GFP^ *(LEU2), mad2Δ::TRP1* | 4E, S1A, S2E |
| 4556 | *MAT*a*, trp1Δ63 ura3-52 his3Δ200 lys2-8Δ1, spc105Δ::NAT, leu2Δ-1::* Spc105 ^222::GFP^ *(LEU2), mad2Δ::TRP1* | 4E |
| 4557 | *MAT*a*, trp1Δ63 ura3-52 his3Δ200 lys2-8Δ1, spc105Δ::NAT, leu2Δ-1::*Spc105 ^222::GFP, 101-104, 340-340::AAAA^ *(LEU2), mad2Δ::TRP1* | 4E, S2E |
| 4558 | *MAT*a *trp1Δ63 ura3-52 his3Δ200 lys2-8Δ1, spc105Δ::NAT, leu2Δ-1::* Spc105 ^222::GFP, 101-104, 340-340::AAAA^ *(LEU2), mad2Δ::TRP1* | 4E |
| 4949 | *MAT*a *trp1Δ63 ura3-52 his3Δ200 lys2-8Δ1, spc105Δ::NAT1, leu2Δ-1::* Spc105 ^222::GFP, RASA (V76, F78::A)^ *(LEU2), mad2Δ::TRP1* | 4E |
| 4950 | *MATα, trp1Δ63 ura3-52 his3Δ200 lys2-8Δ1, spc105Δ::NAT1, leu2Δ-1::* Spc105 ^222::GFP, RASA (V76, F78::A)^ *(LEU2), mad2Δ::TRP1* | 4E |
| 4951 | *leu2Δ-1 trp1Δ63 ura3-52 his3Δ200 lys2-8Δ1, mad2Δ::TRP1* | 4E |
| 5007 | *MATα, leu2Δ-1 trp1Δ63 ura3-52 his3Δ200 lys2-8Δ1, bub3Δ::KAN* | 4E |
| 5009 | *MAT*a*, trp1Δ63 leu2Δ-1 his3Δ200 lys2-8Δ1, spc105Δ::NAT1, ura3-52::* Spc105 ^222::GFP, RASA (V76, F78::A)^ *(URA3), bub3Δ::KAN* | 4E |
| 5010 | *MAT*a*, trp1Δ63 leu2Δ-1 his3Δ200 lys2-8Δ1, spc105Δ::NAT1, ura3-52::* Spc105 ^222::GFP, RASA (V76, F78::A)^ *(URA3), bub3Δ::KAN* | 4E |
| 4682 | *MAT*a*, trp1Δ63 leu2Δ-1 his3Δ200 lys2-8Δ1, spc105Δ::NAT, Spc105^222::GFP^ (LEU2), bub3Δ::KAN* | 4E |
| 4683 | *MAT*a*, trp1Δ63 leu2Δ-1 his3Δ200 lys2-8Δ1, spc105Δ::NAT, Spc105^222::GFP^ (LEU2), bub3Δ::KAN* | 4E |
| 4737 | *trp1Δ63 leu2Δ-1 his3Δ200 lys2-8Δ1, spc105∆::NAT,Spc105 ^101-104, 340-343::AAAA 222::GFP^ (LEU2), bub3∆::KAN* | 4E |
| 4738 | *trp1Δ63 leu2Δ-1 his3Δ200 lys2-8Δ1, spc105∆::NAT,Spc105 ^101-104, 340-343::AAAA 222::GFP^ (LEU2), bub3∆::KAN* | 4E |
| 5699 | *MATα, trp1Δ63 leu2Δ-1 his3Δ200 lys2-8Δ1, bub1ΔK-mCherry-HYG, mad2Δ::TRP1* | 4F |
| 5635 | *MATa, trp1Δ63 leu2Δ-1 his3Δ200 lys2-8Δ1, spc105Δ::NAT, Spc105 ^101-104, 340-343::AAAA 222::GFP^ (LEU2), mad2Δ::TRP1, bub1ΔK-mCherry-HYG* | 4F |
| 5636 | *MATa, trp1Δ63 leu2Δ-1 his3Δ200 lys2-8Δ1, spc105Δ::NAT, Spc105 ^101-104, 340-343::AAAA 222::GFP^ (LEU2), mad2Δ::TRP1, bub1ΔK-mCherry-HYG* | 4F |
| 5637 | *MATa, trp1Δ63 leu2Δ-1 his3Δ200 lys2-8Δ1, spc105Δ::NAT, Spc105 ^RASA (V76, F78::A) 222::GFP^ (LEU2), mad2Δ::TRP1, bub1ΔK-mCherry-HYG* | 4F |
| 5638 | *MATa, trp1Δ63 leu2Δ-1 his3Δ200 lys2-8Δ1, spc105Δ::NAT, Spc105 ^RASA (V76, F78::A) 222::GFP^ (LEU2), mad2Δ::TRP1, bub1ΔK-mCherry-HYG* | 4F |
| 4979 | *trp1Δ63 leu2Δ-1 ura3-52 his3Δ200 lys2-8Δ1, mad2Δ::TRP1, sgo1Δ::KAN* | 4F |
| 4980 | *MATα, trp1Δ63 leu2Δ-1 ura3-52 his3Δ200 lys2-8Δ1, mad2Δ::TRP1, sgo1Δ::KAN* | 4F |
| 4981 | *MATα, trp1Δ63 ura3-52 his3Δ200 lys2-8Δ1, spc105Δ::NAT1, leu2Δ-1::* Spc105 ^222::GFP, RASA (V76, F78::A)^ *(LEU2), mad2Δ::TRP1, sgo1Δ::KAN* | 4F |
| 4982 | *MATα, trp1Δ63 ura3-52 his3Δ200 lys2-8Δ1, spc105Δ::NAT1, leu2Δ-1::* Spc105 ^222::GFP, RASA (V76, F78::A)^ *(LEU2), mad2Δ::TRP1, sgo1Δ::KAN* | 4F |
| 4978 | *MATα, trp1Δ63 ura3-52 his3Δ200 lys2-8Δ1, spc105Δ::NAT1, leu2Δ-1::* Spc105 ^222::GFP, RASA (V76, F78::A)^ *(LEU2), mad2Δ::TRP1, sgo1Δ::KAN* | 4F |
| 5698 | *MATa, trp1Δ63 ura3-52 his3Δ200 lys2-8Δ1, fpr1Δ, FRB-GFP-Spc105 (LEU2), Spc72-mCherry-HYG, Glc7-1XFKBP12-HIS3, prMet3-Cdc20 (URA3)* | 5A, S3B |
| 5495 | *MAT*a, *trp1Δ63 ura3-52 his3Δ200 lys2-8Δ1, leu2Δ-1::* FRB-GFP-Spc105 *(LEU2), Glc7-1XFKBP-HIS3, SPC72-*mCherry*-HYG, prMET3-CDC20 (URA3)* | 5B |
| 4733 | *MATa, trp1Δ63 ura3-52 his3Δ200 lys2-8Δ1,* *spc105Δ::NAT, Spc105^222::GFP 105-107::EEE, S109E^ (LEU2), Bub1-ymCherry-Hyg* | 5C |
| 4734 | *MATa, trp1Δ63 ura3-52 his3Δ200 lys2-8Δ1,* *spc105Δ::NAT, Spc105^222::GFP 105-107::EEE, S109E^ (LEU2), Bub1-ymCherry-Hyg* | 5C |
| 4778 | *MAT*a *, trp1Δ63 ura3-52 his3Δ200 lys2-8Δ1,* *spc105Δ::NAT, leu2Δ-1::* Spc105^222::GFP, 101-104::AAAA^ *(LEU2), sgo1Δ::Kan* | 5D |
| 5087 | *MAT*a*, trp1Δ63 ura3-52 his3Δ200 lys2-8Δ1, spc105Δ::NAT1, leu2Δ-1::* Spc105 ^222::GFP, 105-107::EEE, S109E^ *(LEU2), sgo1Δ::KAN* | 5D |
| 5088 | *MAT*a*, trp1Δ63 ura3-52 his3Δ200 lys2-8Δ1, spc105Δ::NAT1, leu2Δ-1::* Spc105 ^222::GFP, 105-107::EEE, S109E^ *(LEU2), sgo1Δ::KAN* | 5D |
| 5548 | *MATa, trp1Δ63 ura3-52 his3Δ200 lys2-8Δ1, spc105Δ::NAT, Spc105^222::GFP 105-107::EEE, S109E^ (LEU2), dam1Δ::TRP1::dam1(S257A S265A S292A)::KAN-MX* | 5D |
| 5549 | *MATa, trp1Δ63 ura3-52 his3Δ200 lys2-8Δ1, spc105Δ::NAT, Spc105^222::GFP 105-107::EEE, S109E^ (LEU2), dam1Δ::TRP1::dam1(S257A S265A S292A)::KAN-MX* | 5D |
| 5186 | *MAT*a*, trp1Δ63 ura3-52 his3Δ200 lys2-8Δ1, spc105Δ::NAT1, leu2Δ-1::* Spc105 ^222::GFP, 101-104::AAAA^ *(LEU2), sgo1Δ::KAN* | 5D, S2C |
| 5187 | *MATα, trp1Δ63 ura3-52 his3Δ200 lys2-8Δ1, spc105Δ::NAT1, leu2Δ-1::* Spc105 ^222::GFP, 101-104::AAAA^ *(LEU2), sgo1Δ::KAN* | 5D |
| 5188 | *MAT*a*, trp1Δ63 ura3-52 his3Δ200 lys2-8Δ1, spc105Δ::NAT1, leu2Δ-1::* Spc105 ^222::GFP, S77D^ *(LEU2), sgo1Δ::KAN* | 5D |
| 5162 | *MAT*a *trp1Δ63 ura3-52 his3Δ200 lys2-8Δ1, spc105Δ::NAT1, leu2Δ-1::* Spc105 ^222::GFP, S77D, 105-107::EEE, S109E^ *(LEU2), sgo1Δ::KAN* | 5D |
| 5163 | *MAT*a*, trp1Δ63 ura3-52 his3Δ200 lys2-8Δ1, spc105Δ::NAT1, leu2Δ-1::* Spc105 ^222::GFP, S77D, 105-107::EEE, S109E^ *(LEU2), sgo1Δ::KAN* | 5D |
| 3442 | *MAT*a *trp1Δ63 ura3-52 his3Δ200 lys2-8Δ1, spc105Δ::NAT, leu2Δ-1::* Spc105 ^222::GFP^*^,^* ^101-104::AAAA^ *(LEU2)* | S1A |
| 4526 | *MAT*a*, trp1Δ63 ura3-52 his3Δ200 lys2-8Δ1, spc105Δ::NAT, leu2Δ-1::* spc105 ^222::mCherry, 101-104, 340-340::AAAA^ *(LEU2), SGO1-*GFP*-HIS3* | S2A |
| 4527 | *MAT*a*, trp1Δ63 ura3-52 his3Δ200 lys2-8Δ1,spc105Δ::NAT, leu2Δ-1::* Spc105 ^222::mCherry^*(LEU2), SGO1-*GFP*-HIS3* | S2A |
| 4528 | *MAT*a*, trp1Δ63 ura3-52 his3Δ200 lys2-8Δ1, spc105Δ::NAT, leu2Δ-1::* Spc105 ^222::mCherry^*(LEU2), SGO1-*GFP*-HIS3* | S2A |
| 5086 | *MAT*a *, spc105Δ::NAT, leu2Δ-1::* Spc105 ^222::GFP, 340-343::AAAA^ *(LEU2), sgo1Δ::KAN* | S2C |
| 5381 | *MAT*a*, trp1Δ63 ura3-52 his3Δ200 lys2-8Δ1, spc105Δ::NAT, leu2Δ-1:: BP (*Spc105^79-145^)-Spc105 ^222::GFP, 101-104, 340-340::AAAA^ *(LEU2),* bub1ΔK*-mCherry-HYG* | S2D |
| 5382 | *MAT*a*, trp1Δ63 ura3-52 his3Δ200 lys2-8Δ1, spc105Δ::NAT, leu2Δ-1:: BP (*Spc105^79-145^)-Spc105 ^222::GFP, 101-104, 340-340::AAAA^ *(LEU2),* bub1ΔK*-mCherry-HYG* | S2D |
| 5389 | *MAT*a*, trp1Δ63 ura3-52 his3Δ200 lys2-8Δ1, spc105Δ::NAT, leu2Δ-1:: RVSF+BP (*Spc105^55-145^)-Spc105 ^222::GFP, 101-104, 340-340::AAAA^ *(LEU2),* bub1ΔK*-mCherry-HYG* | S2D |
| 5390 | *MAT*a*, trp1Δ63 ura3-52 his3Δ200 lys2-8Δ1, spc105Δ::NAT, leu2Δ-1:: RVSF+BP (*Spc105^55-145^)-Spc105 ^222::GFP, 101-104, 340-340::AAAA^ *(LEU2),* bub1ΔK*-mCherry-HYG* | S2D |
| 5749 | *MATa, trp1Δ63 ura3-52 his3Δ200 lys2-8Δ1, spc105Δ::NAT, RVSF (*Spc105^55-145, 101-104::AAAA^)*+Spc105 222::GFP 101-104::AAAA (LEU2),bub1ΔK-mCherry-HYG* | S2D |
| 5750 | *MATa, trp1Δ63 ura3-52 his3Δ200 lys2-8Δ1, spc105Δ::NAT, RVSF (*Spc105^55-145, 101-104::AAAA^)*+Spc105 222::GFP 101-104::AAAA (LEU2),bub1ΔK-mCherry-HYG* | S2D |
| 5505 | *MATa, trp1Δ63 ura3-52 his3Δ200 lys2-8Δ1, fpr1Δ, FRB-GFP-Spc105 (LEU2), Glc7-1XFKBP12-HIS3, Bub3-ymCherry-HYG* | S3A |
| 5506 | *MATa, trp1Δ63 ura3-52 his3Δ200 lys2-8Δ1, fpr1Δ, FRB-GFP-Spc105 (LEU2), Glc7-1XFKBP12-HIS3, Bub3-ymCherry-HYG* | S3A |
| 5817 | *MATa, trp1Δ63 ura3-52 his3Δ200 leu2∆-1, can1, Nsl1-GFP-NAT, Spc72-mCherry-HYG* | S3C |
| 5818 | *MATa, trp1Δ63 ura3-52 his3Δ200 leu2∆-1, can1, Nsl1-GFP-NAT, Spc72-mCherry-HYG* | S3C |
| 5819 | *MATa, trp1Δ63 ura3-52 his3Δ200 leu2∆-1, can1, Nsl1-GFP-NAT, Spc72-mCherry-HYG, lys2::LYS2-GAL-HO Glc7-spc105 ^RASA (V76, F78::A)^* | S3C |
| 5820 | *MATa, trp1Δ63 ura3-52 his3Δ200 leu2∆-1, can1, Nsl1-GFP-NAT, Spc72-mCherry-HYG, lys2::LYS2-GAL-HO Glc7-spc105 ^RASA (V76, F78::A)^* | S3C |
| 3380 | *MATa, trp1Δ63 ura3-52 his3Δ200 leu2∆-1, can1* | S3C |
| 2675 | *MATa, trp1Δ63 ura3-52 his3Δ200 leu2∆-1, can1, lys2::LYS2-GAL-HO Glc7-spc105 ^RASA (V76, F78::A)^* | S3C |
