## Supplementary material for "Minimization of a harmful cross-talk between mitotic checkpoint silencing and error correction": Table S2

Table S2: Plasmids used in this study:

| Plasmid | Origin | Parent | Description |
| --- | --- | --- | --- |
| pAJ108 | Joglekar lab | pSK954 | ndc80-6A (T21A, S37A, T54A, T71A, S95A, S100A)-12Myc (*KAN*) |
| pSB148 | Biggins lab | pRS316 | *prIPL1+IPL1+trIPL1 (CEN, URA3)* |
| pSB617 | Biggins lab | - | dam1(S257A S265A S292A), (*KAN)* |
| pAJ449 | This study | pRS305 | *prSPC105*+Spc105^222::GFP^+*trSPC105 (LEU2)* |
| pAJ492 | This study | pRS305 | *prSPC105*+Spc105^222::GFP, 340-343::AAAA^+*trSPC105 (LEU2)* |
| pAJ521 | This study | pET28a+MBP | HIS(X6)-MBP-Spc105^222::GFP^ (1-455) |
| pAJ525 | This study | pRS305 | *prSPC105*+Spc105^222::GFP, 101-104::AAAA^+*trSPC105 (LEU2)* |
| pAJ526 | This study | pRS305 | *prSPC105*+Spc105^222::GFP, 101-104, 340-343::AAAA^+*trSPC105 (LEU2)* |
| pAJ528 | This study | pET28a+MBP | HIS(X6)-MBP-Spc105^222::GFP, 101-104, 340-343::AAAA^ (1-455) |
| pAJ603 | This study | pRS305 | *prSPC105*+FRB-GFP-Spc105+*trSPC105* (*LEU2*) |
| pAJ632 | This study | pRS305 | *prSPC105*+TOG2 (Stu2^311-550^)+ Spc105^222::GFP^+*trSPC105 (LEU2)* |
| pAJ633 | This study | pRS305 | *prSPC105*+TOG2 (Stu2^311-550^)+ Spc105^222::GFP, 101-104::AAAA^ +*trSPC105 (LEU2)* |
| pAJ671 | This study | pRS316 | *prIPL1+ipl1-2* (H352Y)*+trIPL1 (CEN, URA3)* |
| pAJ689 | This study | pRS305 | *prSPC105*+Spc105^79-145^(BP)+Spc105^222::GFP, 101-104::AAAA^ +*trSPC105* *(LEU2)* |
| pAJ695 | This study | pRS305 | *prSPC105*+Spc105^222::mCherry^ +*trSPC105* *(LEU2)* |
| pAJ696 | This study | pRS305 | *prSPC105*+Spc105^222::mCherry, 101,104, 340-343::AAAA^+*trSPC105 (LEU2)* |
| pAJ743 | This study | pRS305 | *prSPC105*+Spc105^222::GFP, 105-107::EEE, S109E^+*trSPC105 (LEU2)* |
| pAJ775 | This study | pRS305 | *prSPC105*+Spc105^222::GFP, RASA (V76, F78::A)^+*trSPC105 (LEU2)* |
| pAJ805 | This study | pRS305 | *prSPC105*+Spc105^222::GFP, S77D^+*trSPC105 (LEU2)* |
| pAJ806 | This study | pRS305 | *prSPC105*+Spc105^222::GFP, S77D, 105-107::EEE, S109E^ +*trSPC105 (LEU2)* |
| pAJ817 | This study | pSK954 | *prSPC105*+Spc105^222::mCherry, 101-104::AAAA,340-343::AAAA^+*trSPC105 (KAN)* |
| pAJ818 | This study | pSK954 | *prSPC105*+Spc105^222::mCherry^+*trSPC105 (KAN)* |
| pAJ840 | This study | pRS305 | *prSPC105*+Spc105^56-145^(RVSF+BP)+Spc105^222::GFP, 101-104::AAAA^ +*trSPC105* *(LEU2)* |
| pAJ885 | This study | pRS305 | *prSPC105*+Spc105^56-145, 101-104::AAAA^ (RVSF)+Spc105^222::GFP, 101-104::AAAA^ +*trSPC105* *(LEU2)* |
| pAJ887 | This study | pRS305 | *prSPC105*+Spc105^222::GFP, GILK::AAAA^ +*trSPC105* *(LEU2)* |
| pAJ890 | This study | pRS305 | *prSPC105*+Spc105^222::GFP, GILK::AAAA,^ ^101-104::AAAA^ +*trSPC105* *(LEU2)* |
